# Neutral genetic diversity during selective sweeps in non-recombining populations

**DOI:** 10.1101/2024.09.12.612756

**Authors:** Sachin Kaushik, Kavita Jain, Parul Johri

## Abstract

Selective sweeps, resulting from the spread of beneficial, neutral, or deleterious mutations through a population, shape patterns of genetic variation at linked neutral sites. While many theoretical, computational, and statistical advances have been made in understanding the genomic signatures of selective sweeps in recombining populations, relatively less is understood in populations with little/no recombination, and arbitrary dominance and inbreeding. Using diffusion theory, we obtain the full expression for the expected site frequency spectrum (SFS) at linked neutral sites immediately post and during the fixation of moderately or strongly beneficial mutations. When a single hard sweep occurs, the SFS decays as 1/*x* for low derived allele frequencies (*x*), similar to the neutral SFS at equilibrium, whereas at higher derived allele frequencies, it follows a 1/*x*^2^ power law as also seen in a rapidly expanding neutral population. We show that these power laws are universal in the sense that they are independent of the dominance and inbreeding coefficients, and also characterize the SFS during the sweep. Additionally, we find that the derived allele frequency where the SFS shifts from the 1/*x* to 1/*x*^2^ power law is inversely proportional to the selection strength; thus under strong selection, the SFS follows the 1/*x*^2^ dependence for most allele frequencies. When clonal interference is pervasive, the SFS immediately post-fixation becomes U-shaped and can be approximated by the equilibrium SFS of selected sites. Our results will be important in developing statistical methods to infer the timing and strength of recent selective sweeps in asexual populations, genomic regions that lack recombination, and clonally propagating tumor populations.

## Introduction

Genetic variation is determined by multiple evolutionary processes such as mutation, recombination, population size changes, and selection. While selection against deleterious mutations can affect genome-wide patterns of variation (Charlesworth, 2013), local effects of fixation of beneficial mutations along the genome, also known as selective sweeps (Stephan, 2019), can be used to identify loci involved in recent positive selection. The effect of fixation of a new beneficial mutation at linked sites was first described by Maynard Smith and Haigh (1974) who obtained analytical expressions for the reduction in nucleotide diversity as a function of the distance from the beneficial mutation. Although Maynard Smith and Haigh restricted their study to describing patterns of diversity due to the fixation of a single allele, numerous studies that followed developed the relevant theory to include the effects of recurrent hitch-hiking, wherein new beneficial mutations arise continually at the relevant mutation rate and could occur at any position along the chromosome. Both coalescence theory (Kaplan et al., 1989) and diffusion equations (Stephan et al., 1992) were used to obtain the expected number of segregating sites and diversity in the presence of recurrent hitchhiking, respectively. Note that all these analyses assumed that only a single beneficial mutation was on its path to fixation at a time, which is a reasonable assumption in highly recombining species such as *Drosophila melanogaster*.

An insight into the skew in the expected site frequency spectrum (SFS) of neutral alleles as a result of genetic hitchhiking due to a beneficial allele was first investigated using coalescent simulations by Braverman et al. (1995) and subsequently, analytical expressions were obtained (Barton, 1998, 2000; Fay and Wu, 2000; Durrett and Schweinsberg, 2004). It was shown that immediately post-fixation of a strongly beneficial mutation, the SFS became U-shaped, with a larger proportion of high frequency and low frequency variants, while a lower proportion of intermediate frequency variants than expected under strict neutrality. As neutral diversity recovered post-fixation due to new mutations, the skew towards the high-frequency alleles returned to equilibrium levels faster than the skew towards low frequency alleles. Thus while the SFS is U-shaped after a single fixation event, it becomes predominantly left-skewed when recurrent hitchhiking is considered, as shown by the exact expressions for the neutral SFS using diffusion theory (Kim, 2006). Although much work has been done to model the effects of selective sweeps in recombining populations, including incorporating background selection as the null model (Kim and Stephan, 2000), accounting for interference between beneficial mutations (Kim and Stephan, 2003), spatial structure in populations (Slatkin and Wiehe, 1998; Barton, 2000), incorporating sweeps from standing variation (Berg and Coop, 2015) and the development of maximum likelihood methods to detect selective sweeps by studying the spatial pattern of diversity and linkage disequilibria that are characteristics of selective sweeps (Kim and Stephan, 2002; Kim and Nielsen, 2004), relatively less is known about sweep signatures in non-recombining populations.

Arguments based on the coalescence theory suggest that in the absence of recombination, a star genealogy will be produced, with new mutations contributing to the low frequency alleles (Figure 2 of Fay and Wu (2000)). Concordantly, coalescent-based simulations with no recombination indicated a prevalence of only low frequency alleles due to hitchhiking (Simonsen et al., 1995; Fu, 1997) with the shape of the SFS resembling that of a neutral logistically growing population (Fu, 1997). The haplotype variation generated by neutral mutations after a recent beneficial fixation in non-recombining populations was characterized by Messer and Neher (2012), who showed how the haplotype frequency spectrum immediately post-fixation is different from that expected under neutral equilibrium. Subsequently, Kosheleva and Desai (2013) obtained expressions for the site frequency spectrum of beneficial and linked neutral mutations in asexual populations. However, the latter study was restricted to scenarios where there is pervasive clonal interference, relevant to large microbial populations. While asymptotic properties of the site frequency spectra in asexual populations have been studied previously, we lack a detailed theory that describes the full SFS due to ongoing and completed selective sweeps in fully non-recombining populations, when selective sweeps are rare. We here use a diffusion theory approach to model selective sweeps in a non-recombining population and provide expressions for the SFS at linked neutral sites for arbitrary dominance and probability of inbreeding. Using extensive forward simulations, we test the accuracy of our theory and show how the shape of the SFS depends on the strength of selection, dominance and inbreeding coefficient, and the frequency of beneficial mutations in the population. Our model is most suitable for species or genomic regions with little or no recombination, such as the Y or W sex chromosomes (Charlesworth, 2017), and clonally reproducing organisms including tumor populations (Durrett, 2015).

## Model

We first consider a haploid asexual or equivalently a fully non-recombining diploid population of constant size *N* with *L* biallelic sites in which all but one site are neutral. Note that our diploid model assumes independent segregation of chromosomes. Thus, a fully clonal diploid population is better modeled as a haploid asexual population with 2*L* biallelic sites. The wildtype and mutant alleles at the neutral sites are denoted, respectively, by *a* and *A*, and at the selected site, by *a*^*∗*^ and *A*^*∗*^, with respective fitnesses, 1 and 1 + *s, s >* 0. It is clear that if only one neutral site is fully linked to the selected locus and the mutant allele at the selected locus eventually fixes, all the individuals will carry the neutral allele that was initially linked to the beneficial mutation thus reducing the neutral diversity to zero (Atwood et al., 1951). For this reason, in the classic literature on selective sweeps in two-locus models (Maynard Smith and Haigh, 1974), one considers a recombining population. Here, in contrast, we consider an infinite-loci model for non-recombining populations in which, as in Kimura’s infinite-sites model (Kimura, 1969, 1971) for fully recombining population, all the *L* → ∞ neutral sites linked to the selected locus are initially fixed for the wildtype allele, and irreversible mutation occurs at neutral sites with rate *µ* per site per generation (see Fig. 1). Thus, while diversity is continually generated at the neutral sites due to mutations, it can be lost due to random genetic drift or effects of selection, resulting in nontrivial patterns of diversity in a non-recombining population, as illustrated in Fig. 1.

**Figure 1.**
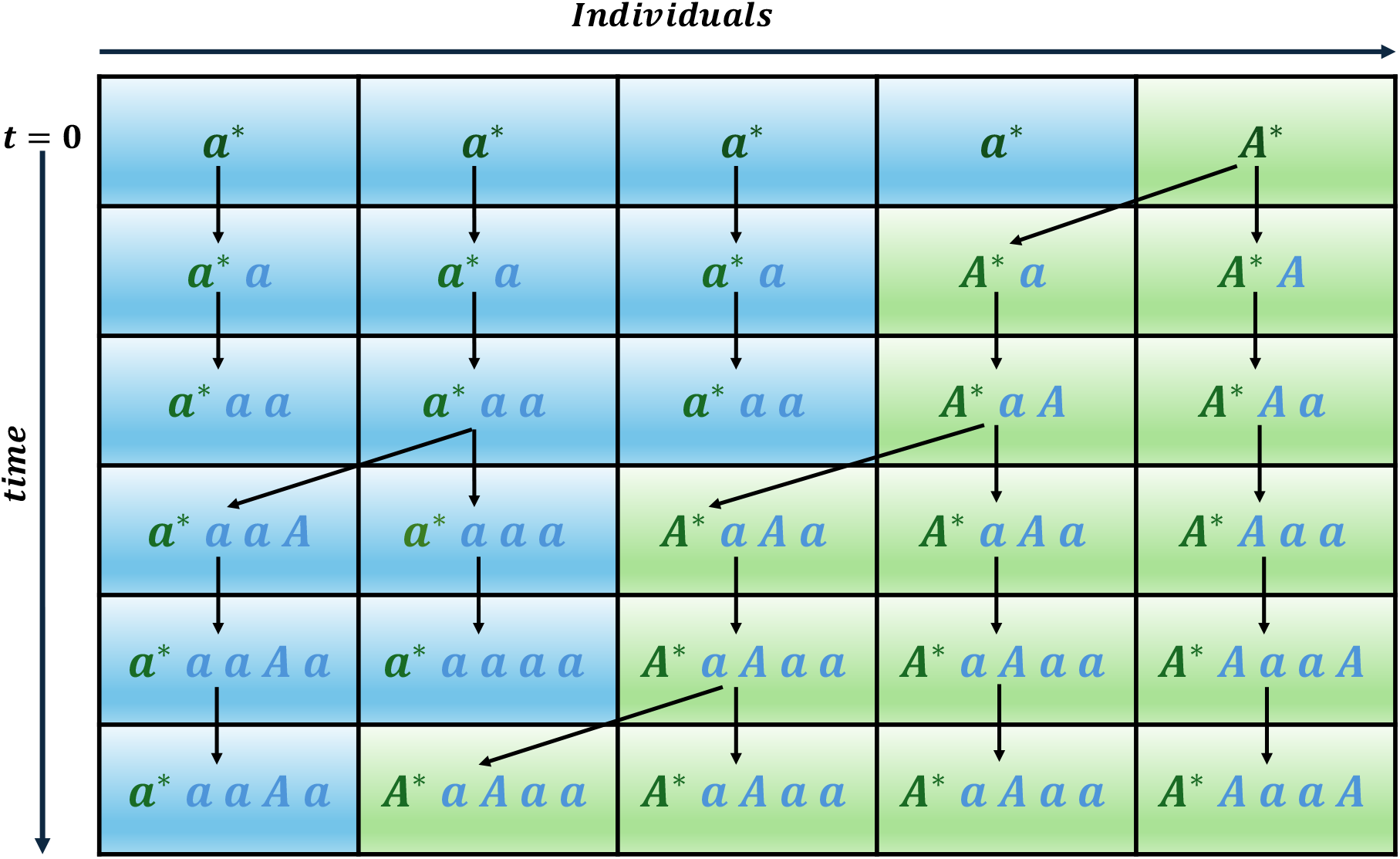
Schematic representation of the model for a haploid asexual population of size *N* = 5. The first locus (where the alleles are marked with a star) represents the selected site, while the remaining sites are neutral. At *t* = 0, all the neutral sites are fixed for the wildtype allele *a* and a single beneficial mutation *A*^*∗*^ arises at the selected site. New neutral mutations, denoted by *A*, arise in subsequent generations according to the infinite-sites model. Here, the selected site and only those neutral sites that are polymorphic are depicted. The population evolves in time following Wright-Fisher dynamics as indicated by the arrows. At the time of sampling, the SFS at the neutral sites is derived from the subpopulation (depicted with a green background) that contains the beneficial mutation at the selected locus. In the example provided, at *t* = 5, the neutral mutant counts across the four neutral sites are {1, 3, 0, 1}, corresponding to frequencies of {1/4, 3/4, 0/4, 1/4}, with 4 being the size of the subpopulation. The SFS is obtained by creating a histogram from these frequencies. The above model is generalized to include dominance, inbreeding and recurrent beneficial mutations; see text for details.

The model described above can be extended in a straightforward manner to include dominance and inbreeding. At the selected locus, we assume that the three genotypes *A*^*∗*^*A*^*∗*^, *a*^*∗*^*A*^*∗*^ and *a*^*∗*^*a*^*∗*^ have fitnesses 1 + *s*, 1 + *hs* and 1, respectively, where 0 *< h <* 1 is the dominance coefficient. Then if *p* and *q* = 1 − *p* denote the deterministic frequency of the *A*^*∗*^ and *a*^*∗*^ allele, respectively, and 0 *< F <* 1 is the inbreeding coefficient, in an inbreeding population, these genotypes occur with frequencies *p*^2^ + *Fpq*, 2*pq*(1 − *F*), and *q*^2^ + *Fpq*. At time *t* = 0, a single copy of the beneficial allele, *A*^*∗*^ is introduced at the selected site, and then the population evolves following the standard Wright-Fisher dynamics in forward time using the genotypic frequencies and fitnesses mentioned above. These dynamics are implemented in numerical simulations as described below in Methods.

In the following, we numerically study how neutral diversity is affected due to a single hard sweep at the selected site during and at the end of the selective sweep when all the sites are fully linked; although we are mainly interested in a fully non-recombining population, we also perform some simulations for weak recombination. As a measure of diversity, we consider the (unfolded) SFS *f* (*x, t*) of the neutral mutant allele with frequency *x* at time *t* in the *subpopulation* carrying the selected allele *A*^*∗*^ as shown in Fig. 1. Since we are interested in the dynamics of the subpopulation carrying the selected allele, one needs to consider only a neutral population whose size is changing with time. Now assuming that the alleles at the neutral sites evolve independently (as in Kimura’s infinite-sites model) but in an expanding population with instantaneous size 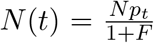, we formulate a diffusion theory for the SFS which is described in Appendix A and Appendix B, and study it in detail in Appendix C and Appendix D. These results are compared with those obtained from extensive simulations of diploid non-recombining populations, and are discussed in Results. Some important extensions of the model described above, *viz*., the effect of clonal interference and deleterious sweeps are also discussed.

## Methods

### Forward time simulations

All forward time simulations are conducted using SLiM version 4.0.1 (Haller and Messer, 2023).

### Single sweep

We simulate a Wright-Fisher population consisting of *N* diploid individuals with no recombination and a mutation rate, *µ* = 10^−7^ per site per generation. Mutations are assumed to be semidominant (*h* = 0.5), partially recessive (*h* = 0.2), or partially dominant (*h* = 0.8), as specified. The simulated genomic region of length *L* = 10^4^ consists of a single selected site where beneficial mutations occur, while the rest of the region is assumed to be completely neutral. For a single selective sweep, a beneficial mutation with selection coefficient, *s >* 0 is introduced at the selected site. The number of sites simulated is chosen such that there is sufficient neutral diversity at the end of the selective sweep. Since the conditional mean fixation time, 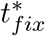 of the mutant allele at the selected locus scales approximately as 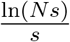 for a haploid population and the mean absorption time of a single neutral mutation, *T*_*abs*_ ∼ ln *N* (Ewens, 2004), neutral diversity at all the *L* sites where a mutant appears at rate *Nµ* per site vanishes if the total absorption time 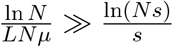. We therefore require 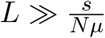 which, for the typical parameters used here, is satisfied.

A single beneficial mutation is introduced at the selected site. If this mutation is lost, the simulation is restarted. This process is repeated until the beneficial mutation fixes in the population. The frequencies of alleles at neutral sites are recorded immediately post-fixation of the selected allele, and the final site frequency spectrum (SFS) in each plot is constructed by averaging over 10^6^ fixation events. Two different initial conditions are tested - (1) when the population has no diversity (i.e., it is monomorphic at all sites), and (2) when the neutral mutations in the population are present at equilibrium frequencies at the time of introduction of the beneficial mutation. The SFS obtained post-fixation is independent of these initial conditions (Fig. S1). While the second initial condition is biologically more realistic, especially when selective sweeps are rare, the first condition is mathematically simpler and is therefore employed throughout this study for obtaining the theoretical results. Correspondingly, all single-sweep simulations shown here are conducted such that the population is initially monomorphic.

The neutral SFS is also obtained when the frequency of the beneficial mutation reaches 0.5 and 0.7 for the first time during its trajectory (conditional on the eventual fixation of the beneficial mutation). Note that the SFS obtained during the fixation of the beneficial mutation refers to the SFS of the neutral alleles only from the subpopulation that carries the beneficial mutation, which becomes equivalent to the entire population after the fixation of the beneficial allele. More specifically, the SFS of neutral alleles is calculated only from the haplotypes that carry the beneficial mutation. The nucleotide site diversity is also calculated for the subpopulation with the beneficial allele at the selected site, at various time points during its path to fixation.

To investigate the effect of inbreeding on the SFS, we extend our simulations to include varying rates of selfing with selfing probabilities *σ*=0 and 0.9. The inbreeding coefficient *F* is related to the selfing probability by the relation 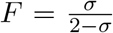 (Wright, 1969) so that *F* = 0 and 0.81, respectively. Simulations of deleterious sweeps (Johri et al., 2021; Kaushik and Jain, 2021) are conducted in a similar fashion but limited to weak selection (|*Ns*| = 2) as the probability of fixation of deleterious mutations decreases exponentially with increasing strength of selection.

### Recurrent sweeps with clonal interference

To understand the effect of clonal interference on the SFS immediately post-sweep, we consider a more complex genomic architecture that includes multiple beneficial mutations arising at selected sites. Two different types of beneficial mutations are simulated: the first type of beneficial mutations (referred to as the *m*_1_ class) have selective effects that follow an exponential distribution and a high beneficial mutation rate (denoted by *µ*_*b*1_), and are expected to cause interference. The second type of beneficial mutations (*m*_2_ class) have fixed selective effects and very low mutation rates (*µ*_*b*2_ → 0) thus minimizing any possibility of interference amongst them; the fixation of beneficial mutations in this class are of interest.

Specifically, in our simulations, the genome architecture consists of a total length of *L* = 11, 000 base pairs with the first *L*_*sel*_ = 10^3^ base pairs consisting of sites where either a neutral or a beneficial mutation from either the first or the second class can occur with mutation rates per site/generation denoted by *µ*_*n*_, *µ*_*b*1_, and, *µ*_*b*2_ respectively. The rest of the genome consists of neutral sites where, as before, the mutation rate per generation is *µ* and at which the SFS is computed. We assume that the total mutation rate at the selected part of the genome is also *µ* with *µ*_*n*_ + *µ*_*b*1_ + *µ*_*b*2_ = *µ* and *µ*_*n*_ = 0.895*µ, µ*_*b*1_ = 0.1*µ, µ*_*b*2_ = 0.005*µ*. The selective effect of mutations in class *m*_1_ are chosen from an exponential distribution with mean *Ns*_1_ = 100 while that of class *m*_2_ mutations is fixed with scaled selection strength *Ns*_2_ = 50. As the mutation rate of *m*_2_ is sufficiently low so that only a single *m*_2_ mutation (“focal mutation”) arises in the population until its fixation, the SFS of linked neutral alleles is obtained immediately after the fixation of the focal beneficial mutation following a burn-in period of 10*N* generations to allow for the background (*m*_1_ class) beneficial mutations and neutral mutations to reach equilibrium frequencies.

### Effect of recombination

Our model assumes no and very low rates of recombination such that it is only applicable to regions with recombination rates that are much lower than the strength of selection of the beneficial allele so that the beneficial mutant fixes before recombination events can occur. More specifically, the model presented here is appropriate when the force of recombination is weak (*Nr «* 1) but selection is strong (*Ns* 1) where *r* is the recombination rate per site/generation. In order to investigate whether our model would remain accurate within these limits, we perform simulations with an extremely low rate of recombination (*Nr* = 10^−6^) and a significantly higher one (*Nr* = 10^−3^).

### Obtaining the SFS and nucleotide diversity

At the end of each fixation event, all or *n*=10 diploid individuals are sampled from the population. At each polymorphic site and for each individual, we record the frequency of neutral mutations and store this information in a vector *v*_1_. We then generate a histogram of the *v*_1_ vector converting allele counts into allele frequencies while maintaining a constant area under the histogram. This procedure is repeated for 10^6^ fixation events, and the averaged histogram is obtained. The resulting averaged data is then binned with a bin size of 100 to obtain the final SFS shown in the figures. The diversity is calculated by averaging 2*x*(1 − *x*) across all genomes, where *x* represents the frequency of the neutral mutation at each neutral site (also see Eq. (9) below). The SFS is obtained both post-fixation and during the selective sweep. While obtaining the SFS during the fixation, i.e., at the intermediate stages, the population consists of two subpopulations: one with the beneficial allele at the selected site and the other with a neutral allele at the selected site. Only the subpopulation that carries the beneficial allele was used to make the SFS.

Python scripts for generating the SFS described above are provided at https://github.com/JohriLab/sweeps_in_nonrecombining_populations. Simulation parameters such as the length of the genomic region, scaled selection coefficient, recombination rates, inbreeding coefficient, population size, and mutation rates of both selected and neutral sites are detailed in figure legends or specified as needed.

### Theoretical expectations from the infinite-sites model

To obtain an insight into the numerical results, we also studied the expected SFS in the framework of diffusion theory.

### Single sweep

As described in Appendix A, the neutral SFS *f* (*x, t*) in a temporally changing population of effective size *N*_*e*_(*t*) can be described by the following partial differential equation (Evans et al., 2007):

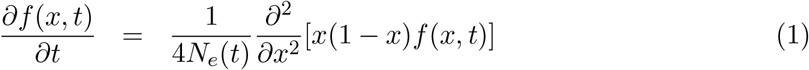

where, as detailed below, 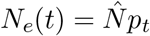 due to changing size of the subpopulation with selected allele and 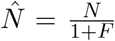 is the effective population size due to inbreeding alone (Pollak, 1987; Glémin, 2012). The partial differential equation, Eq. (1), is subject to boundary conditions (Evans et al., 2007)

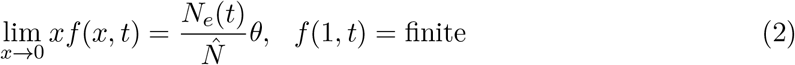

where,

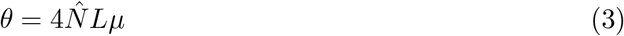

and to ensure equivalence between the single-locus diffusion theory and forward time simulations for *L* sites, the mutation rate in the diffusion model is scaled to *Lµ*. At low allele frequency, Eq. (2) models the balance between the loss of polymorphism due to the absorption of the mutant allele by genetic drift and the gain by new mutations. Furthermore, as there are no polymorphic neutral sites when the selected allele arises, the initial frequency *f* (*x*, 0) = 0.

As discussed in Appendix B, for strong selection (*Ns* 1), the effective population size can be approximated by 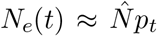 where the mutant allele frequency *p*_*t*_ at the selected locus evolves deterministically according to

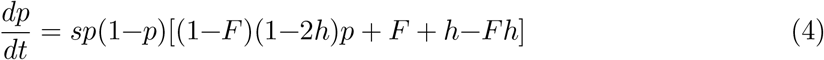

with initial frequency (Maynard Smith, 1976),

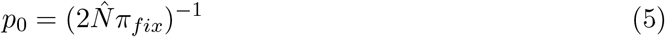

where, the fixation probability of a single beneficial mutant is given by

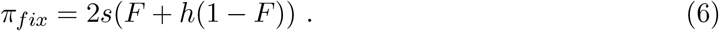

Thus in the framework of diffusion theory, Eq. (1)-Eq. (6) yield the SFS, *f* (*x, t*), at a given time slice.

To find the SFS immediately post-fixation, we require the distribution of time to fixation conditional on fixation, which has been derived for randomly mating population (Martin and Lambert, 2015; Kaushik and Jain, 2021; Götsch and Bürger, 2024), and for an inbred population (Kaushik, 2023) using a semi-deterministic theory. But the expressions for these distributions are approximate and not explicit (barring the case of *h* = 1/2, *F* = 0). Also, as shown in Fig. S2, the SFS calculated by averaging it over the conditional distribution of fixation time is not significantly different from that obtained by using the conditional mean fixation time except when selection is weak and/or the selected allele is recessive. Therefore, here we obtain the SFS immediately post-fixation at the conditional mean fixation time, 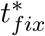 of the selected allele. For a large randomly mating population, 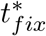 is given by (Ewing et al., 2011)

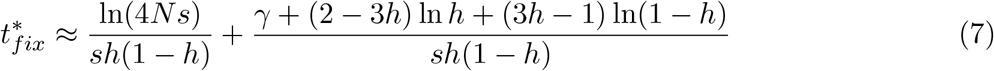

where *γ* is the Euler constant, whereas for an inbred population, it is obtained from equation (17) and (18) of Glémin (2012). Note that for small values of *Ns*, more accurate and explicit expressions for the time to fixation have been derived (Charlesworth, 2020). The time-dependent SFS *f* (*x, t*) is, however, calculated at the time (*t*) when a given value of deterministic allele frequency *p*_*t*_ is reached; this time, *t*, is obtained by integrating Eq. (4).

The normalized sample SFS, *f*_*n,i*_(*t*) in a sample of size *n « N* can be obtained from *f* (*x, t*) through

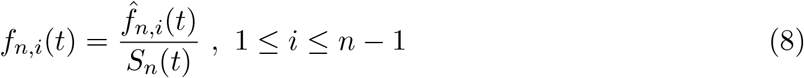

where the unnormalized sample SFS, 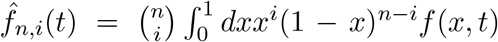 and 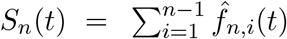 is the mean number of segregating sites in the sample of size *n* at time *t*. The mean diversity is then obtained from the unnormalized sample SFS as

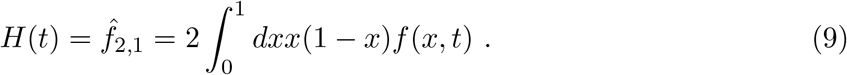

Although we do not provide analytical expressions for these reduced quantities, they are obtained using the numerical solution of Eq. (1) in the above definitions.

In Appendix C, the exact solution of Eq. (1) is described and an approximate analytical expression for the time-dependent SFS is obtained in Appendix D (see Eq. (13)and Eq. (14) below). These analytical results as well as the exact numerical solution of Eq. (1) is compared with the simulations in the Results section. As it is easier to integrate Eq. (1) numerically instead of computing the sum in its exact solution given by Eq. (C.2), the numerical solution of the diffusion equation is used throughout the article which is obtained using Mathematica 13.1.1, and is available in a Mathematica file at https://github.com/JohriLab/sweeps_in_nonrecombining_populations.

### Clonal interference and multiple mutations

When multiple beneficial mutations arise in a non-recombining population, competition between beneficial mutations can result in a decrease in their probability of fixation and is referred to as clonal interference (Gerrish and Lenski, 1998). We assume that a focal beneficial mutation with selective advantage *s*_2_ and dominance coefficient *h*_2_ experiences clonal interference from other background beneficial mutations with selective advantage *s*_1_ and dominance coefficient *h*_1_. Then the reduced fixation probability of the beneficial mutation at the focal site is given by (Gerrish and Lenski, 1998),

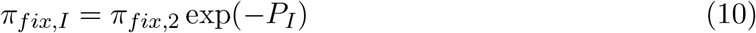

where the expected number of interfering mutations are

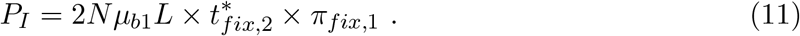

In the above expressions, *π*_*fix,i*_ and 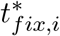 for *i* = 1, 2 are obtained by replacing *s* and *h* in Eq. (6) and Eq. (7) by *s*_*i*_ and *h*_*i*_, respectively. The SFS at the neutral sites is then obtained using *π*_*fix,I*_ instead of *π*_*fix*_ in Eq. (5).

As we discuss later, when the rate of new beneficial mutations is high enough so that *P*_*I*_ *>* 1, the bulk of the SFS at neutral sites immediately post fixation of the beneficial mutation at the focal site can be described by the equilibrium SFS of selected sites, *f*_*sel*_(*x*), under the infinite-sites model with selection coefficient *s*_1_ and dominance coefficient *h*_1_. In the absence of inbreeding, this is given by (Jain and Kaushik, 2022)

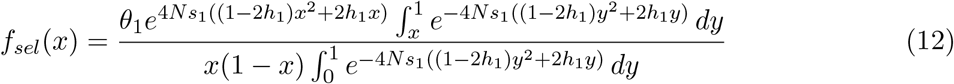

Where 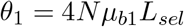. As the selective effect of mutations in the class *m*_1_ is chosen from an exponential distribution, the above SFS is averaged over this distribution to obtain the final equilibrium SFS, *(f*_*sel*_(*x*)).

## Data availability

All scripts to perform simulations, numerical computations, and calculation of statistics have been made available at https://github.com/JohriLab/sweeps_in_nonrecombining_populations.

## Results

### Single hard sweep

The site frequency spectrum (SFS) of linked neutral alleles when a single hard sweep occurs at the selected locus is shown for various parameters in Figs. 2-5, and below, we discuss how the SFS depends on them. Note that the theoretical results obtained here represent the expected SFS and the corresponding SFSs obtained via simulations represent an averaged value over 10^6^ fixations for each scenario. To understand the amount of variability across independent simulation replicates, see Fig. S3 and the Discussion section.

#### Crossover in the distribution of allele frequencies

In Fig. 2, the simulation data for the SFS at the completion of the sweep shows that below a crossover allele frequency *x*_*c*_, the SFS decreases as *x*^−1^ - this is expected as the alleles at low frequencies behave in a neutral fashion and the equilibrium SFS in a large neutral population, 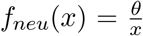 (Wright, 1938; Kimura, 1968). But for higher frequencies, the SFS decays as *x*^−2^ which has been observed in exponentially growing populations (Durrett, 2013, 2015). Figs. 3-5 show the simulation data for the ratio 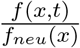: for *x « x*_*c*_, we find that this ratio is constant with respect to the allele frequency which means that the SFS decays as *x*^−1^, at the same rate as the neutral expectation, but for *x x*_*c*_, it decreases with the allele frequency so that the SFS *f* (*x, t*) observed during and post-sweep, decreases faster than the neutral expectation. The inset of Fig. 2 shows the normalized sample SFS, which is defined in Eq. (8), and reinforces the conclusion that the fraction of polymorphic sites at the end of selective sweep relative to that for neutral equilibrium SFS is negligible for large *i*, where *i* is the derived allele count in a sample SFS (Braverman et al., 1995; Fay and Wu, 2000).

To understand these results quantitatively, in Appendix D, we show that when the selective sweep is nearly complete, the time-dependent SFS can be approximated by

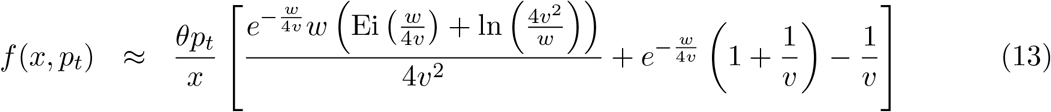

Where 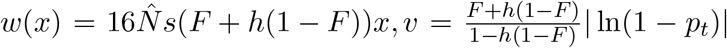, and Ei(*z*) is the exponential integral function. On expanding the RHS of Eq. (13)when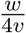 is smaller and larger than one, we find that

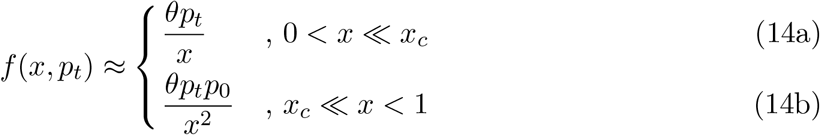

where the scaled mutation rate *θ* is given by Eq. (3), the frequency *p*_*t*_ of the selected allele at time *t* is determined by Eq. (4), the initial frequency *p*_0_ is given by Eq. (5) and the crossover from the *x*^−1^ to *x*^−2^ behavior occurs at an allele frequency, *x*_*c*_, determined by setting 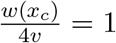. *x*_*c*_ is given by

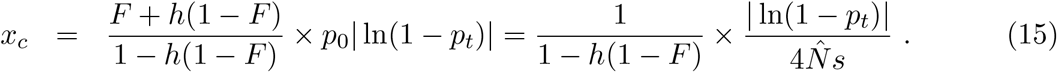

Thus Eq. (14b) shows that the decay exponent two of the SFS at large frequencies is *universal* in the sense that it is independent of the dominance and inbreeding coefficients, and holds even before the completion of the sweep. But the crossover allele frequency *x*_*c*_ given by Eq. (15) depends on various parameters of the model.

Figure 2 shows a comparison between the approximate solution Eq. (13)and the asymptotic results in Eq. (14) when they are evaluated at the mean fixation time 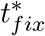 given by Eq. (7), and we find that they agree well with those obtained from the stochastic forward-in-time simulations for the asexual population.

**Figure 2.**
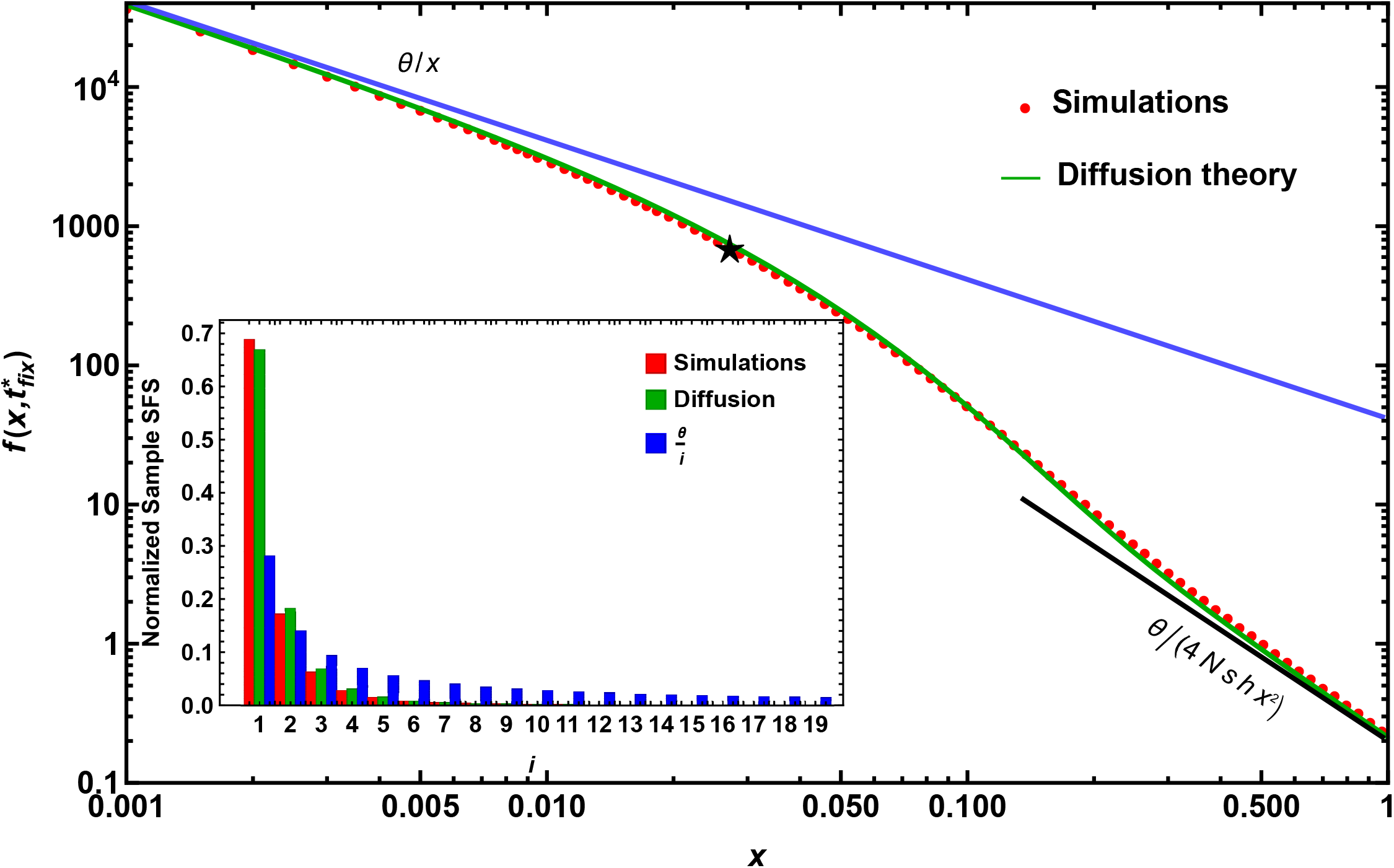
The site frequency spectrum (SFS), i.e., the expected distribution of allele frequencies at neutral sites fully linked to a single new beneficial mutation immediately post-fixation. Here, the red points represent results obtained from forward time simulations, while the green line corresponds to the analytical expression in Eq. (13)which is obtained using diffusion theory. The blue and black solid lines, respectively, depict the result in Eq. (14a) for the SFS at neutral equilibrium and Eq. (14b) for higher derived allele frequencies on using *p*_*t*_ = 0.996, *h* = 1/2, *F* = 0 in these expressions. The crossover frequency *x*_*c*_ is obtained from Eq. (15) and is indicated by a black star. The inset shows the normalized SFS from a sample of 20 haploid genomes obtained from simulations and diffusion theory, with the neutral equilibrium sample SFS shown for comparison. Here, the parameters used are *N* = 10^3^, *µ* = 10^−7^, *L* = 10^5^, *s* = 0.1, *h* = 0.5, *F* = 0.

#### Strength of selection

Figure 3 shows the SFS immediately post-fixation for three values of the scaled selection parameter *Ns* (keeping *N* fixed); these results for a diploid non-recombining (or haploid asexual) population are also compared with those obtained from diffusion theory by numerically integrating Eq. (1). We find that the proportion of moderate to high frequency alleles is consistently smaller in simulations than that obtained from theory, and the quantitative match between the results for the two models is relatively poor for 0.01 *< x <* 0.1; presently we do not have an understanding of this discrepancy.

**Figure 3.**
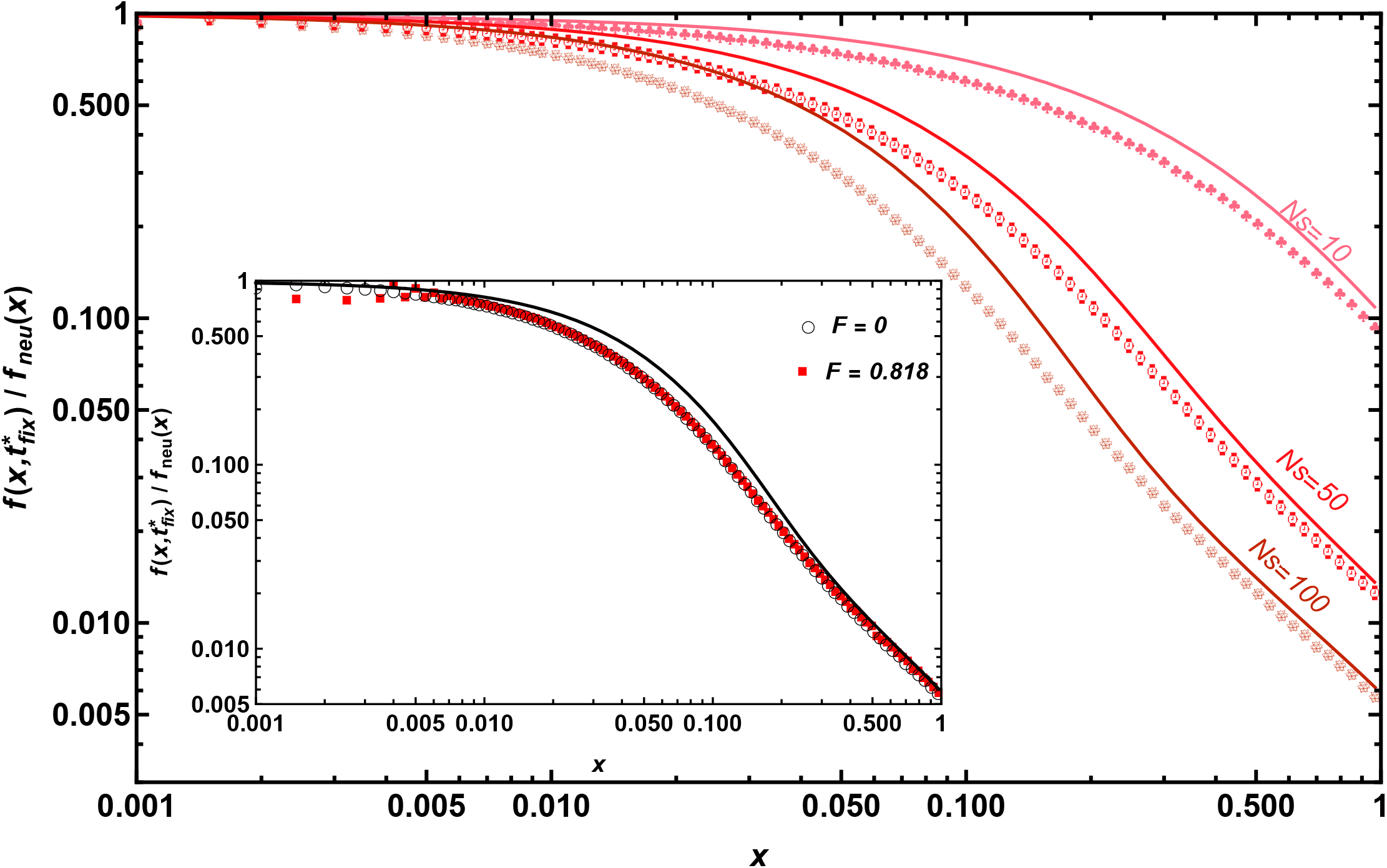
The SFS at neutral sites fully linked to a single new beneficial mutation immediately post-fixation relative to the neutral equilibrium SFS, *f*_*neu*_(*x*) = *θ*/*x*, for varying strengths of selection: *Ns* = 10, 50, 100 (top to bottom). Here, the points are obtained from forward-in-time simulations and the corresponding solid lines are obtained by numerically solving Eq. (1) for the SFS using diffusion theory. The inset shows the relative SFS for varying inbreeding coefficients. Again, points are obtained from forward-in-time simulations whereas the black line is obtained using diffusion theory described by Eq. (1). The parameters used are *N* = 10^3^, *µ* = 10^−7^, *L* = 10^5^, *F* = 0 (main), *s* = 0.1 (inset), and *h* = 0.5.

However, the qualitative conclusions drawn from our diffusion theory on the dependence of the SFS on the strength of selection are in agreement with the simulation results. We find that as expected, selection strength weakly affects the SFS at small allele frequencies. But, as Fig. 3 shows, the expected number of polymorphic sites at high allele frequencies is greater for smaller selection coefficients. This observation is consistent with the theoretical prediction in Eq. (14b) which shows that the prevalence of high-frequency derived alleles is inversely proportional to the initial frequency, *p*_0_ which is proportional to (4*Ns*)^−1^ due to Eq. (5) and Eq. (6). Note that because the population is fully non-recombining and the SFS is from neutral alleles in the *subpopulation*, high-frequency derived alleles consist only of mutations that occurred in the early stages of the selective sweep, while low-frequency derived alleles comprise mutations that occurred during the later stages of the sweep, i.e. closer to the time near fixation. When selection on the selected allele is stronger, the initial frequency of the selected allele is small but the deterministic growth in its frequency occurs soon after it arises and it fixes quickly in the population. Therefore, there are fewer derived alleles that hitch-hike with the beneficial allele to high frequencies. On the other hand, when selection is weaker, the frequency of the beneficial allele after which it is likely to increase deterministically is much higher (Kaplan et al., 1989; Kim and Stephan, 2002). Moreover, as the fixation time of the selected allele is longer, more haplotypes will go to fixation. Thus, there will be more high-frequency derived alleles when selection is weak.

Equation (15) suggests that the crossover allele frequency, *x*_*c*_, decreases with increasing *Ns*. This is because *x*_*c*_ is inversely proportional to the factor *Ns*, while directly proportional to the logarithmic factor 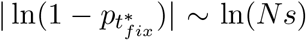 since 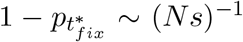when selection is strong and the initial frequency is small. The decrease in *x*_*c*_ with *Ns* is consistent with the simulation data shown in Fig. 3 as the ratio 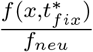 remains close to one for a larger range of allele frequencies when selection is weaker. This also implies that under very strong selection, the SFS follows a *x*^−2^ behavior for most of the allele frequency range, with the window of allele frequencies exhibiting the *x*^−1^-dependence being significantly reduced.

In Fig. 3, the selection coefficient, *s*, is varied while keeping the population size constant. From Eq. (14b), for a given selection coefficient, the product *θp*_0_ is independent of *N*, and thus the tail of the SFS is independent of the population size. Indeed, the simulation data in Fig. S4 for the SFS immediately post-fixation supports this theoretical expectation for allele frequency, *x >* 0.5.

#### Inbreeding

Here, we relax the assumption of random mating and investigate the effect of inbreeding on the SFS immediately post-fixation. For various inbreeding coefficients under semidominance, the data in Fig. S5 indicates that for the entire range of derived allele frequency, the expected number of polymorphic sites decreases with increasing inbreeding coefficient, but the ratio 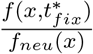 shown in the inset of Fig. 3 is, in fact, independent of the inbreeding coefficient.

These trends are well captured by diffusion theory: for semidominance, as *w*(*x*) and *v* defined by Eq. (13)are independent of *F* while the scaled mutation rate *θ* in Eq. (3) is proportional to (1 + *F*)^−1^, it follows from Eq. (13)that *f* (*x, t*) decreases with increasing *F* but the ratio 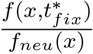 is unaffected by inbreeding. Moreover, Eq. (5) suggests that for arbitrary dominance coefficients, *p*_0_ is only mildly different when inbreeding is high. As a result, the neutral SFS at higher derived allele frequencies immediately post-sweep is less impacted by the dominance coefficient in a highly inbred population as shown in Fig. S6.

#### Dominance

In Fig. 4, we compare the SFS at the end of the selective sweep when the selected allele is partially recessive, semidominant, and partially dominant for a fixed selection strength and no inbreeding. The simulation data shows that when the derived allele frequencies are small, the SFS is independent of the dominance coefficient; for moderately large *x*, the expected number of polymorphic sites are higher for higher dominance coefficient but this trend reverses at higher derived allele frequencies. We also note that the number of polymorphic sites observed in the simulated non-recombining population is consistently smaller than the corresponding results from diffusion theory. This discrepancy is most prominent for the partially recessive allele and for moderate allele frequencies in the range 10^−2^ − 10^−1^. Due to this mismatch between results from diffusion theory and simulations, the dependence of the crossover frequency on the dominance coefficient as predicted by Eq. (15) also does not align with the trends observed from simulations.

**Figure 4.**
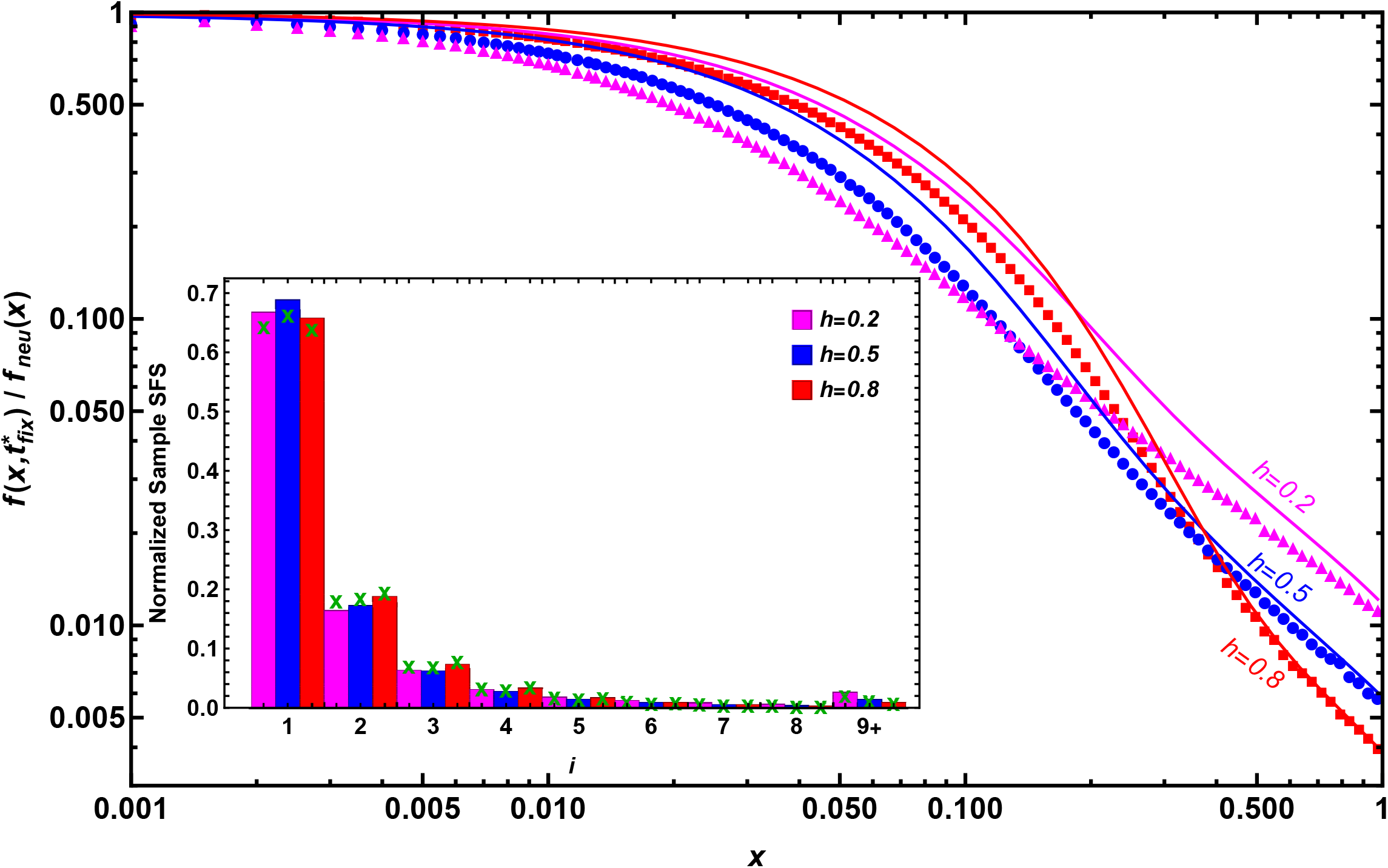
The SFS at neutral sites fully linked to a single new beneficial mutation immediately post-fixation relative to the neutral equilibrium SFS, *f*_*neu*_(*x*) = *θ*/*x*, for various dominance coefficients: *h* = 0.2, 0.5, 0.8. Here, the points are obtained from forward time simulations whereas the corresponding lines of the same color are obtained using diffusion theory by numerically solving Eq. (1). The inset shows the normalized sample SFS for various dominance coefficients where the bars and crosses show the data obtained from simulations and diffusion theory respectively. Note that in the normalized sample SFS shown here, the frequency of derived alleles for counts larger than 8 (i.e., *i* ≥ 9) have been pooled together. The other parameters used are *N* = 10^3^, *µ* = 10^−7^, *s* = 0.1, *F* = 0, and *L* = 10^5^.

However, the dominance patterns in the tail of the SFS are captured by Eq. (14b) which shows that for *F* = 0, the SFS for *x x*_*c*_ is inversely proportional to the dominance coefficient. This is consistent with the simulation results in Fig. 4 where an increase (decrease) in the proportion of derived high frequency alleles is observed when the beneficial mutation is recessive (dominant) relative to the semidominant case. This result can be understood as follows: for strong selection, as the frequency of the selected allele grows as *e*^*hst*^ during the early phase, the frequency of the beneficial mutation will remain quite small for a longer time period when the mutation is recessive (*h <* 0.5) vs when it is dominant (*h >* 0.5). Consequently, derived neutral alleles are more likely to hitchhike to higher frequencies with the recessive beneficial allele compared to the dominant one, resulting in higher expected number of polymorphic sites for *h* = 0.2 relative to that for *h* = 0.8 in Fig. 4. Whereas for low derived allele frequencies, which are largely produced by mutations that occur close to the completion of the sweep, there are more such alleles when the selected allele is dominant vs recessive. This is due to the longer time to fixation of the dominant allele after reaching a high frequency in the population compared to the recessive one near the end of the sweep. However, note that this effect completely disappears in the sample SFS (refer to the inset in Fig. 4), as it is difficult to observe alleles with frequencies as low as 0.005 in a sample of 20 genomes.

#### Time-dependence of SFS

In the preceding subsections, we discussed the SFS immediately post selective sweep demonstrating that the SFS has a 1/*x*^2^ dependence on higher derived allele frequencies. We now extend our investigation to the neutral SFS *during* the fixation of a beneficial mutation. When the sweep at the selected locus is incomplete, the population consists of two subpopulations: (i) individuals with the beneficial mutation at the selected locus, and (ii) individuals with the wildtype or neutral mutation at the selected locus; here, we focus on the former subpopulation. Fig. 5 shows the simulation data for the time-dependent SFS, *f* (*x, t*), at two time slices where the frequency of the selected allele is far smaller than one. A comparison with the diffusion theory shows that it is not quantitatively accurate when the derived allele frequency is small. Also, as our approximate expression Eq. (14) is not valid for a low frequency of the selected allele, here we discuss the results obtained from simulations and numerically integrating the diffusion equation, Eq. (1).

**Figure 5.**
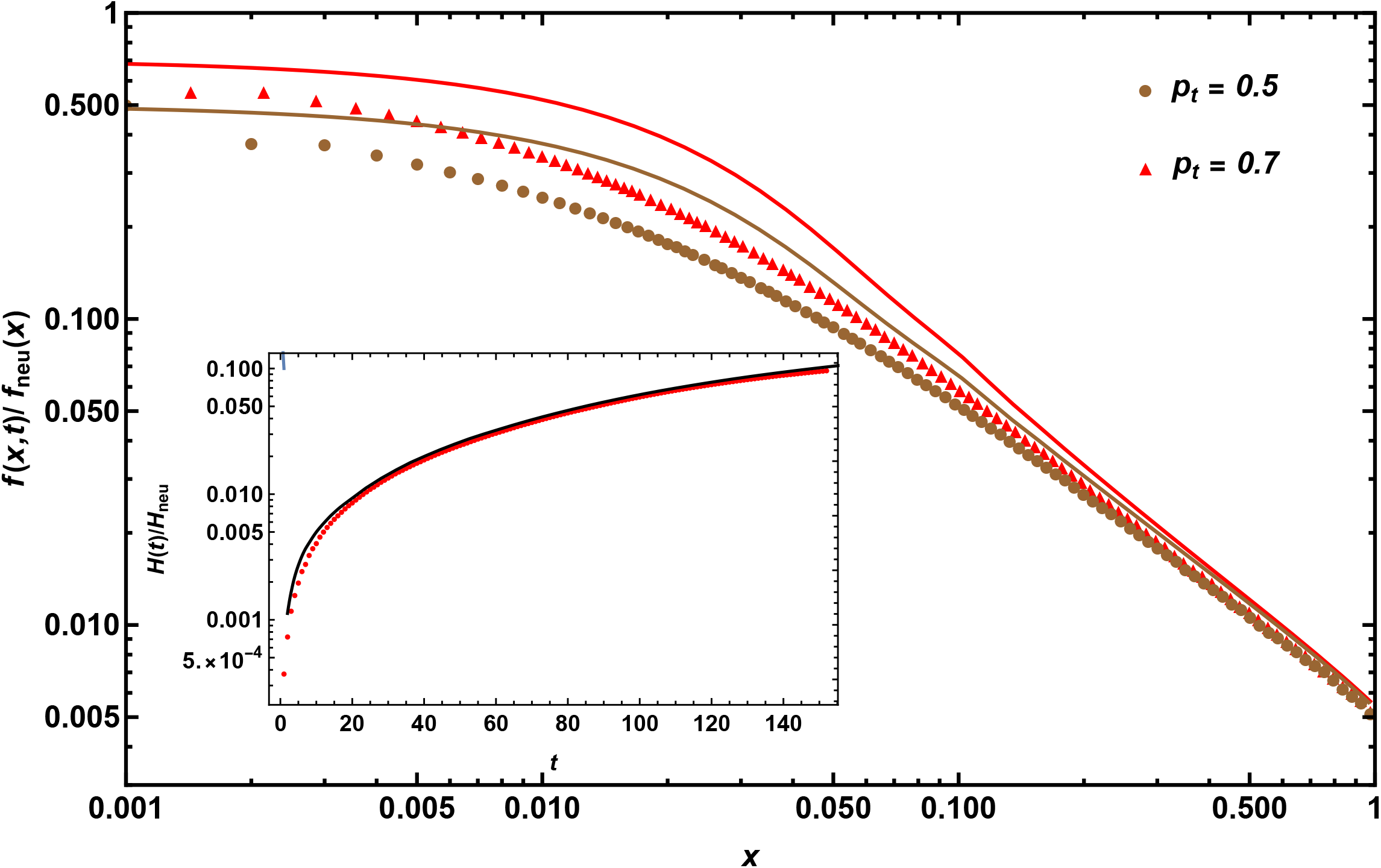
The time-dependent SFS at neutral sites fully linked to a single new beneficial mutation relative to the neutral equilibrium SFS, *f*_*neu*_(*x*) = *θ*/*x* when the beneficial mutation’s frequency *p*_*t*_ is smaller than one. The parameters used for the main figure are *N* = 10^3^, *µ* = 10^−7^, *s* = 0.1, *F* = 0, *h* = 0.5, and *L* = 10^7^. The inset shows the time evolution of neutral nucleotide diversity (*H*(*t*)) in the subpopulation carrying the beneficial mutation, relative to the neutral equilibrium diversity (*H*_*neu*_ = 4*NµL*), during the progression of the beneficial mutation to fixation. The parameters used in the inset are *N* = 500, *µ* = 10^−4^, *h* = 0.5, *F* = 0, and *s* = 0.05. In both the figures, the points are obtained using forward time simulations (brown circles for *p*_*t*_ = 0.5 and red triangles for *p*_*t*_ = 0.7), and the solid lines are obtained using the diffusion equations by numerically solving Eq. (1).

We find that the SFSs obtained from the subpopulation at different time points differ mostly in the number of low-frequency derived alleles with almost no difference in the number of high-frequency derived alleles, with a larger number of low-frequency derived alleles when the sweep is closer to completion. Intuitively, if one samples a sweep closer to fixation, there are many more genomes sampled in the subpopulation (mimicking a scenario where a population has undergone more generations of exponential growth) and thus there are more new mutations that could have occurred recently; this scenario therefore yields many more singletons or low-frequency derived alleles. On the other hand, most high-frequency derived alleles arose very early on, during the beginning of the sweep, and their frequency is independent of the time of sampling the subpopulation (as *r* = 0). Note that this result is specific to the way we are calculating the SFS only in the subpopulation. For the entire population, the proportion of high-frequency derived alleles would likely not be independent of the time of sampling the sweep. In summary, the contribution of time since the beneficial mutation arose is restricted to low-frequency derived alleles and remains visible in the unnormalized sample SFS although this effect disappears in the normalized sample SFS, as illustrated in Fig. S7.

The dynamics of nucleotide diversity in the subpopulation carrying the beneficial mutation during the course of the sweep (inset of Fig. 5) agrees well with those obtained by solving Eq. (1) numerically except in the early phases of the sweep. This suggests that at least for some statistics (such as nucleotide diversity), diffusion theory is a good approximation even when the selective sweep is far from complete.

#### Recombination

So far, we have discussed the results for a fully non-recombining population where all the neutral sites and the selected locus always remain completely linked, and found that diffusion theory for the infinite-sites model matches well with simulation results for the non-recombining population. Our approach is also applicable to regions with extremely low rates of recombination, when the force of recombination is much weaker than that of selection. These expectations are tested in Fig. 6 which shows that when the recombination rate is extremely small, much smaller than the selection coefficient, the SFS matches the results obtained from diffusion theory. However, when the recombination rate is comparable to the selection coefficient, neutral alleles linked to the selected site can escape fixation by recombining with other backgrounds (individuals with neutral alleles at the selected site), weakening the hitchhiking effect and the frequency spectrum approaches that predicted for recombining populations (Fay and Wu, 2000).

**Figure 6.**
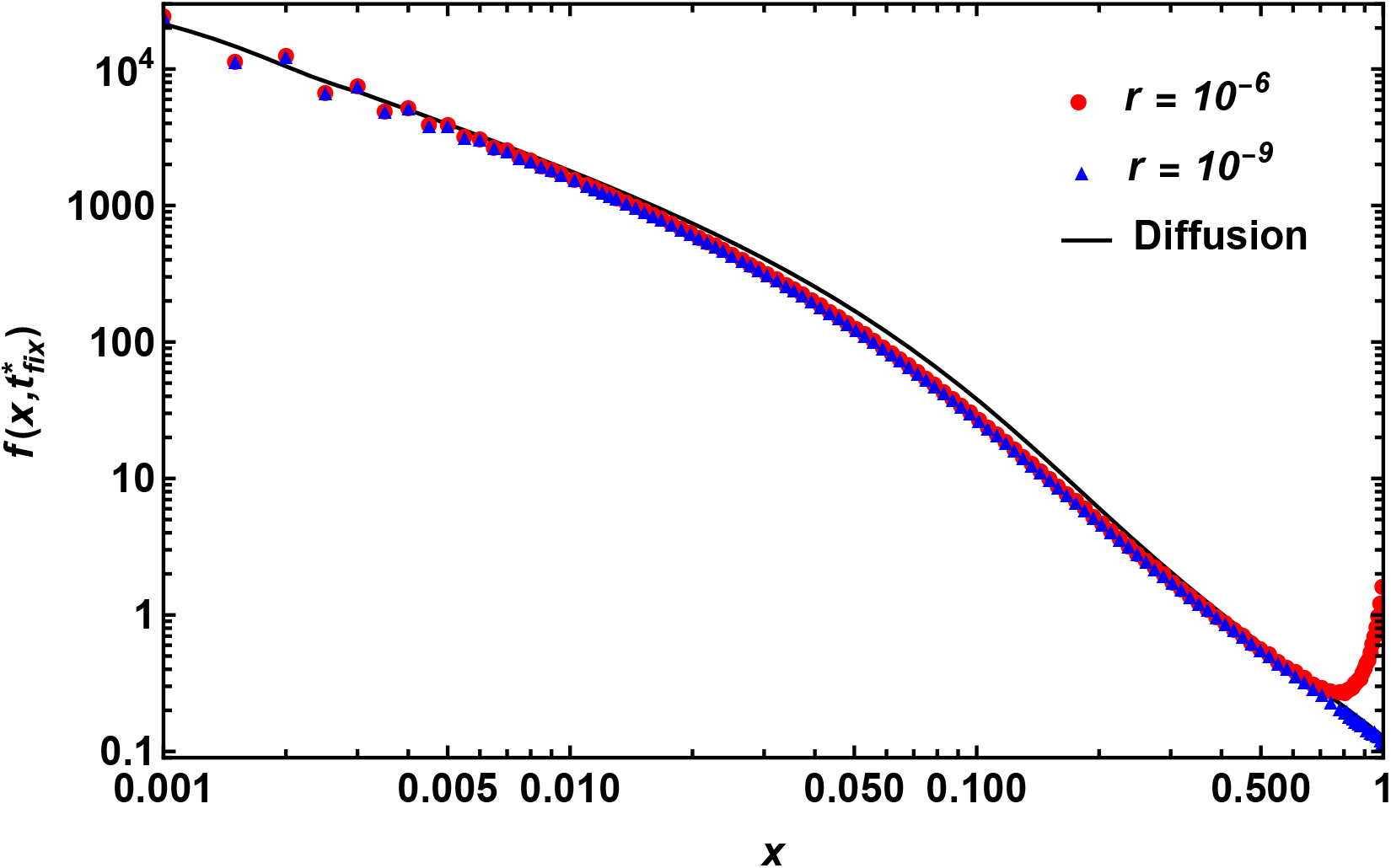
The SFS at neutral sites linked to a single beneficial mutation immediately post-fixation when recombination occurs at scaled rate *Nr* = 10^−3^ (red circles) and 10^−6^ (blue triangles). Points are obtained from forward time simulations while the solid line shows the numerical solution of Eq. (1) for the SFS from diffusion theory. Here parameters used are *N* = 10^3^, *µ* = 10^−6^, *s* = 0.1, *L* = 10^4^, *F* = 0.818, and *h* = 0.5.

To explore the effect of recombination between neutral sites only, we also simulate a neutral population whose size evolves according to the mean frequency of a beneficial mutation conditional on its eventual fixation, and compute the SFS when the population size is 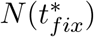 where 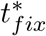 is the conditional mean fixation time given by Eq. (7). Figure S8 demonstrates that the SFS for a neutral population with logistic growth is unchanged in the presence of recombination, confirming that linkage between neutral sites does not influence the SFS post-fixation.

### Clonal interference and multiple mutations

In the previous sections, we have considered a single hard sweep affecting linked neutral genetic variation. However, when multiple beneficial mutations at selected sites occur during the fixation of a single beneficial mutation (here referred to as the “focal” mutation) (Kim and Stephan, 2003; Chevin et al., 2008; Kosheleva and Desai, 2013), the dynamics are more complex, and interference between selected alleles is highly likely in non-recombining populations. Below we explore three different evolutionary regimes with varying degrees of interference amongst selected sites. Note that all the scenarios considered here are characterized by strong selection but the extent of interference is modulated by the rate of new recurrent beneficial mutations entering the population.

In Fig. 7, the simulation data depicts the SFS obtained immediately after the fixation of the focal beneficial mutation (denoted by *m*_2_) which experiences interference from other beneficial mutations (denoted by *m*_1_) which have an exponential distribution of fitness effects. The degree of interference is determined by the expected number of interfering beneficial mutations *P*_*I*_ while the focal mutation is undergoing fixation and is given by Eq. (11). A value of *P*_*I*_ close to or greater than one is likely to result in substantial interference, while a value much smaller than one might indicate almost no interference. In all the cases considered below, the probability of a second *m*_2_ mutation occurring in the population before the fixation of the former *m*_2_ mutation is very low as the expected number of interfering beneficial mutations of type *m*_2_ is 0.07 on using Eq. (11), which is quite low.

**Figure 7.**
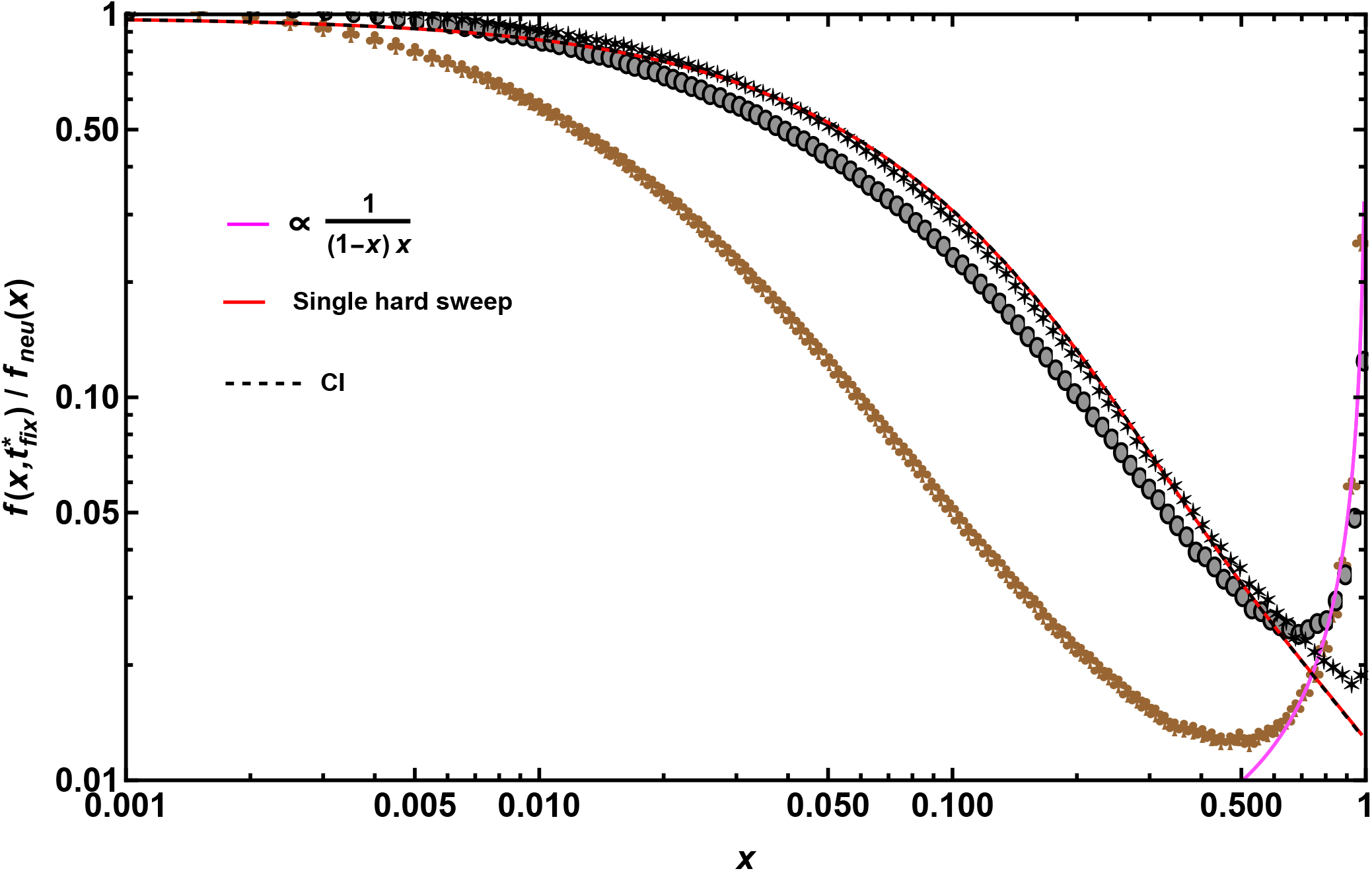
The SFS at linked neutral sites immediately post-fixation of a beneficial allele with *Ns*_2_ = 50 experiencing clonal interference due to strongly beneficial mutations whose distribution is assumed to be exponential with mean *Ns*_1_ = 100. The SFS is shown relative to the neutral equilibrium SFS when the expected number of interfering mutations obtained using Eq. (11) is low (*P*_*I*_ = 0.00184, ***∗***), moderate (*P*_*I*_ = 0.184, grey •) and high (*P*_*I*_ = 1.84, brown *♣*). The points are obtained from forward time simulations, while the red and black dashed lines, respectively, are obtained by numerically solving Eq. (1) assuming no clonal interference and with clonal interference (that is, using Eq. (10) instead of Eq. (6) in the diffusion theory). The magenta line is obtained by integrating Eq. (12) for the equilibrium SFS for selected sites over the exponential distribution of fitness effects of new beneficial mutations. The parameters are *N* = 10^3^, *L* = 11, 000, *µ* = 10^−6^, *F* = 0 and *h*_1_ = *h*_2_ = 0.5.

In the first regime, which we refer to as the “very weak mutation” regime, the rate of new beneficial mutations in *m*_1_ class is kept sufficiently low so that interference between selected alleles is unlikely with *P*_*I*_ ≈ 0.00184. Consequently, as expected, the simulation data for the neutral SFS post-fixation matches our theory for a single selective sweep quite well (shown in Fig. 7) except for derived allele frequencies close to one. As an aside, the match in predictions from simulations and theory in this regime confirms that the expected SFS is independent of the initial condition of the population (see Fig. S1). In the second “weak mutation” regime, the rate of new beneficial mutations was moderately high such that *P*_*I*_ ≈ 0.184. Here, while high frequency derived alleles continue to converge to 1/*x*^2^ for frequencies in the range 0.5 − 0.7, the neutral SFS shows an uptick (when *x >* 0.7), thus forming a U-shaped SFS. In the third “strong mutation” regime characterized by a high rate of new beneficial mutations, interference between beneficial mutations was highly likely (*P*_*I*_ ≈ 1.84) and the effect of selective sweeps at neutral sites is dominated by the presence of multiple beneficial mutations at the selected sites. This intensifies the U-shaped pattern in the SFS, and as shown in Fig. 7, the SFS for high frequency alleles is no longer explained by the 1/*x*^2^ power law.

When the rate of beneficial mutations is sufficiently high and segregating beneficial mutations can interfere with each other, there are two events that determine their dynamics in the population: if new beneficial mutations arise on different haplotypes, they will compete with each other, thus reducing each other’s probability of fixation. On the other hand, if multiple beneficial mutations arise on the same haplotype, then their probability of fixation might be increased. While the former event is likely with moderate levels of interference, the latter becomes more likely as the rate of new beneficial mutations increases and thus the dynamics become dominated by multiple mutations. Frequent hitchhiking due to multiple beneficial mutations has been shown to generate a U-shaped SFS (Kim, 2006) because beneficial alleles at selected sites reach high frequencies and linked neutral alleles hitchhike along with them leading to more number of expected polymorphic sites. Concordantly, we observe a more exaggerated U-shape as the rate of beneficial mutations increases in the population. Therefore, using the single-locus theory described by Eq. (12) for direct selection and assuming that selected alleles follow an exponential distribution of fitness effects, we calculate the expected SFS at equilibrium, *(f*_*sel*_(*x*)) (shown by magenta line in Fig. 7). Interestingly, we find that the expected allele frequency distribution matches our observed SFS from simulations quite well, suggesting that when the extent of clonal interference is high, the SFS at neutral sites is well explained by those at selected sites.

Finally, although the uptick in the high frequency derived alleles is explained well by Eq. (12), we asked if correcting for the probability of fixation with interference using *π*_*fix,I*_ given by Eq. (10) in the diffusion theory would help explain the shape of the observed SFS at the intermediate frequencies. However it does not capture the qualitative shape of the SFS in the intermediate frequencies as shown in Fig. 7. This is expected because our theory does not account for the effect of multiple mutations as this scenario involves multiple subpopulations with varying numbers of beneficial mutations, making it challenging to incorporate clonal interference into the existing framework. In summary, while parts of the SFS post-selective sweep when interference levels are moderate can be captured by the theory presented here, our theory cannot explain the shape of the SFS when interference is common.

### Deleterious sweeps

In this section, we briefly investigate the SFS at linked neutral sites when a single mildly deleterious mutation goes to fixation. In a constant environment, the conditional mean fixation time for a semidominant allele is the same for a beneficial and deleterious mutation of the same magnitude of selection coefficient (Maruyama and Kimura, 1974; Johri et al., 2021; Kaushik and Jain, 2021) although the sojourn time or the equilibrium SFS for a selected allele is not the same for positively or negatively selected mutant allele (Jain and Kaushik, 2022). In our simulations under semidominance (Fig. S6), the neutral SFS after a deleterious mutation fixes is extremely similar to that of a beneficial mutation with the same magnitude of selection coefficient. This result is consistent with previous observations of semidominant mutations in recombining populations (Johri et al., 2021): the post-fixation SFS for a mildly deleterious mutation is similarly skewed as that for a beneficial one. Although simulation data for the semidominant case aligns well with the Maruyama-Kimura symmetry of fixation times (Maruyama and Kimura, 1974), dominance plays a substantial role in determining the trajectory of deleterious fixations (Charlesworth, 2022a). A full theoretical treatment for a deleterious sweep, including an analytical proof for this observed symmetry of the SFS remains an interesting open problem.

## Discussion

In the simplest setting of a panmictic, randomly mating, neutral population of constant size *N* in which a wild type allele mutates at rate *µ* per generation to a mutant allele, the unfolded equilibrium site frequency spectrum (SFS) decays algebraically, as 4*Nµ*/*x* (Wright, 1938; Kimura, 1968; Fisher, 1930a). But this equilibrium result is known to be affected by several factors, such as background selection, selective sweeps, and fluctuations in population size (Fay and Wu, 2000; Przeworski, 2002; Kim and Stephan, 2003; Kim, 2006; Evans et al., 2007; Chevin and Hospital, 2008b; Coop and Ralph, 2012; Neher and Hallatschek, 2013; Kosheleva and Desai, 2013; Charlesworth and Jain, 2014; Živković et al., 2015; Cvijović et al., 2018; Johri et al., 2020; Charlesworth and Jensen, 2021; Jain and Kaushik, 2022; Min et al., 2022; Kaushik, 2023; Balick, 2023). In this study, we focus on the effect of selective sweeps on the SFS at linked neutral sites in species or genomes with little or no recombination. Using extensive numerical simulations and diffusion theory, we study the time-dependent and stationary state SFS, and find that a crossover occurs from the standard neutral theory result of 1/*x* at low allele frequencies to 1/*x*^2^ at high allele frequencies. Although the behavior of the SFS in the latter regime has also been observed in some previous studies on semi-dominant, randomly mating populations (Kosheleva and Desai, 2013; Durrett, 2015), it is not obvious if the *x*^−2^ dependence holds if these assumptions are violated. Here, using a diffusion theory approach, we derive an expression for the time-dependent SFS given by Eq. (13), which shows that these exponents are universal in the sense that they do not depend on the dominance and inbreeding coefficients and are independent of the initial conditions. However, the crossover derived allele frequency given by Eq. (15) depends on the population-genetic parameters in the model; in particular, the frequency at which the crossover occurs is inversely proportional to 4*Ns* suggesting that larger the difference in fitness between the favorable and wildtype genotype, the smaller the frequency at which this crossover occurs. This means that if selection is strong, the majority of the SFS post-fixation will decay as 1/*x*^2^ whereas the transition between the two power laws will only be observed during the fixation of a weakly selected mutation. In fact, the smaller the sample size, the harder it will be to observe the crossover and thus only 1/*x*^2^ is likely to be detected in data.

In models rooted in population genetics, other measures of neutral diversity in expanding populations have also been considered. Specifically, haplotype frequency spectrum (which can be interpreted as an SFS under the infinite-alleles model) was studied by Messer and Neher (2012) using deterministic arguments. Its shape was shown to transition from an exponential function in an asexual population of constant size (assuming strict neutrality) to a power law spectrum when the selective sweep occurs. However, this measure seems difficult to use in practice because it can be challenging to identify the different haplotypes and is usually methodologically limited to the identification of a maximum of 5 or so haplotypes (Nkhoma et al., 2020). This is especially the case when the total number of expected haplotypes is not known (Zhu et al., 2018), as in pathogenic species, when a sufficiently large number of genomes are not sequenced (Booker et al., 2017), long-read data is not available, or single-cell sequencing is not possible. In contrast, allele frequencies are not only easy to get an accurate measure of but also commonly measured in tumor as well as malaria and virus populations, making our framework very generally applicable. Moreover, the shape of the SFS is commonly of theoretical interest amongst population geneticists, as it can be used to predict which evolutionary processes are responsible for evolutionary dynamics in a population, for instance, to assess the rate of adaptation, or to fit historical population size changes. Our approach will, therefore, be important to understand the prevalence of such processes in natural population data of species with little/no recombination as well.

### Comparison to a growing tumor or an expanding neutral population

Motivated by the growth of cancerous cells (Durrett, 2015; Tung and Durrett, 2021; Gunnarsson et al., 2021) and bacterial growth in Luria-Delbrück’s experiments (Cheek and Antal, 2018), recent studies have focused on understanding how neutral diversity is affected in an exponentially growing population using a branching process and shown a similar 1/*x*^2^ decay of the SFS. This behavior can be understood using a simple, purely deterministic argument as follows (Williams *et al*., 2016; Tung and Durrett, 2021): suppose a tumor cell divides at rate *λ*, no death events occur, and mutations occur at rate *µ* per cell per generation with each mutation resulting in a new type (or allele). Then one can ask: how many types of cell are present with a given frequency *x* at an observation time *T* ? Since the population size grows exponentially, the total number of cells at time *T* are *e*^*λT*^, and therefore, a cell type created at time *T* has a frequency, *x* = *e*^−*λT*^. As the number of cell types that have frequency *x* or larger at *T* must have been created before time *T*, adding the number of cell types created until time *T* gives 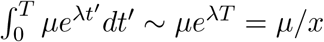. That is, the number of cell types with frequency *x* at *T* is 1/*x*^2^.

If now death events are also allowed so that the population can go extinct with a finite probability, a change from 1/*x* to 1/*x*^2^ behavior occurs at a crossover frequency that increases with the extinction probability (Figure 7a of Gunnarsson et al. (2021)). This behavior is analogous to the crossover seen here as the extinction probability in cancer modeling is analogous to the population size of the Wright-Fisher process, and one may interpret the low (high) cell viability as small (large) population size. Gunnarsson et al. (2021) suggest that this transition represents the transition from a purely birth-death process early on during tumor growth to almost a stable population size in the later stages. This would also be consistent with what we observe in our model of logistic growth, where early on there is exponential growth and later stages mimic a nearly constant-sized population. However, in our study we observe a transition between the two power-laws even when the beneficial mutation has not reached fixation, which would be closer to an exponential and not a logistic growth.

So it is unclear why we see the transition of power laws in our subpopulation before the event of fixation. Note though that the smaller the frequency of the beneficial mutation at the time of sampling (i.e., the further it is from fixation), the smaller the population frequency *x*_*c*_ at which crossover occurs (see 15), and closer is the SFS to following mostly a 1/*x*^2^ power law. This is thus consistent with the argument presented in Gunnarsson et al. (2021). Moreover, this behavior is also consistent with the fact that studies that focussed on modeling only an exponential growth under neutrality (Durrett, 2015) do not observe the crossover, and the stationary state SFS is found to decay as *x*^−2^ for all mutant frequencies in these studies.

Note that we and many previous studies assume a lack of recombination in our models. However, the 1/*x*^2^ power law should remain applicable to an expanding population under strict neutrality even in the presence of recombination (Fig. S8). This is because the SFS measures the *mean* number of polymorphic sites with frequency in the range (*x, x* + *dx*). In fact, it has been shown that the mean number of segregating sites without and with recombination agree when the scaled mutation rate is small enough. But other quantities such as the probability that the population is monomorphic for high mutation rates (Watterson, 1975), or the joint SFS for multiple linked neutral sites (Ferretti et al., 2018) can be substantially different.

### Comparison to selective sweeps in a recombining population

In contrast to the 1/*x*^2^ power law followed in a non-recombining population at large allele frequencies, the SFS immediately post-fixation of a beneficial mutation in a recombining population changes in a non-monotonic fashion at intermediate frequencies (see Equation (5) of Min et al. (2022)) and is a constant at high frequencies (Fay and Wu, 2000; Kim, 2006). Thus while the SFS post selective sweep in a recombining population has high frequency alleles, the SFS in a non-recombining population is only skewed towards low frequency alleles, consistent with Braverman et al. (1995).

Although here we have mostly focused on the SFS during and at the completion of the sweep, we expect that it would recover over time scales of order *N*. Assuming that no more beneficial mutations occur on this time scale, the SFS immediately post-fixation will equilibrate to *f*_*neu*_(*x*) = *θ*/*x*, and it is of interest to find if some range of frequencies take shorter or longer to reach the neutral SFS value (Przeworski, 2002). Our numerical analysis of Eq. (1) shown in Fig. S10 suggests that high frequencies take longer than low and intermediate frequencies to recover; this may be expected as the SFS immediately post-fixation at low frequencies is already close to the neutral equilibrium result. Interestingly, in recombining populations, high frequency alleles recover quickly post-fixation while low frequency alleles take longer to return to equilibrium values. This is also because in a non-recombining population there are no haplotypes that escape during the sweep via recombination (which are responsible for the high frequency alleles observed with recombination). Thus the recovery of high frequency alleles in a non-recombining population must occur gradually as new neutral mutations drift to higher frequencies.

Further work in understanding the recovery of the SFS as well as obtaining the SFS of the entire population is needed, so that expectations under recurrent hitchhiking due to beneficial mutations in non-recombining populations can be obtained. Such efforts can help predict neutral genomic diversity in species with little/no recombination and perhaps better resolve Lewontin’s paradox of variation (Charlesworth and Jensen, 2022). However, other constantly operating evolutionary processes such as background selection will also cause the equilibrium SFS to be strongly left-skewed in a non-recombining population (Cvijović et al., 2018), making the pattern generated by a recent selective sweep quite difficult to distinguish from that of background selection.

Our findings are consistent with previous studies that have evaluated the effect of dominance in generating signals of selective sweeps in recombining populations. In a randomly mating population with recombination, the effect of dominance on reduction in neutral genetic diversity following a hard sweep has been investigated using the mean fixation time of the beneficial allele (Teshima and Przeworski, 2006; Ewing et al., 2011). Recently, in a more general model, the effect of both dominance and inbreeding on the sample SFS following a hard and soft sweep was studied, mainly numerically (Hartfield and Bataillon, 2020), and it was found that these have a relatively weak effect on the sample SFS, which agrees qualitatively with our results in the inset of Fig. 3 and Fig. 4.

### Applicability under pervasive clonal interference

In this study, we find that when the rate of new beneficial mutations is high, i.e., there are multiple beneficial mutations interfering with each other in a non-recombining population, the equilibrium SFS of neutral linked alleles in the population is U-shaped and likely dominated by recurrent the hitchiking effect caused by favorable mutations. In this regime, the SFS post-fixation of beneficial mutations is also U-shaped and no longer follows the 1/*x*^2^ power law. As expected, accounting for the altered fixation probability of beneficial mutations in this regime does not lead to explaining the SFS accurately. This suggests that when clonal interference is pervasive due to a high rate of input of beneficial mutations, the signatures of selective sweeps are not captured by the theory presented here. The neutral (and beneficial) SFS in an asexual population in the clonal interference regime was previously studied in Kosheleva and Desai (2013). In the limit when adaptation occurs very rapidly, an explicit expression for the neutral SFS was obtained which predicts that the SFS decays as *x*^−2^ for small *x* and as ((1 − *x*) ln(1 − *x*))^−1^ for *x* close to one. In this study, we have given a simple argument to understand the nonmonotonic behavior of the SFS at high derived allele frequency which, however, does not capture the ln(1 − *x*) factor mentioned above. We also mention that the U-shaped SFS has also been observed when there is background selection (Cvijović et al., 2018); a comprehensive study incorporating the effect of both positive and negative selection on linked neutral sites is highly desirable.

### Prospects for performing evolutionary inference

The theoretical development presented here is restricted to modeling the subpopulation of haplotypes that carry the beneficial allele. Thus, while we have a general expression for the distribution of allele frequencies at any time, during the progression and post-fixation of the selective sweep, only the scenario post-fixation is comparable to previous theory. It remains a challenge to obtain the SFS of alleles in the entire population, during the progression of the sweep. An expression of the SFS of the entire population would allow us to accurately model partial sweeps and obtain the expected distribution of allele frequencies due to recurrent hitchhiking in completely non-recombining populations. Further work in this direction is needed.

Despite the above caveat, the theory presented here could be useful for estimating the strength of selection and/or time to the origin of a beneficial allele in non-recombining populations, especially in cases where the beneficial variant is known. This will be particularly applicable to sex chromosomes that lack recombination like the human Y chromosome, the 4th or dot chromosome in *Drosophila melanogaster*, haploid asexual organisms or viruses that do not recombine, and can be extended to tumor populations. As propagation of cancerous cells in a tumor involves clonal reproduction and no recombination, the theory presented here could be used to estimate the strength of selection acting on and/or the time of origin of driver mutations (if the driver mutations are known). However, note that our model is applicable for two scenarios without recombination - (1) a haploid case where only a single chromosome is relevant, and (2) a diploid case where there are two chromosomes in each individual that can segregate independently each generation. The second scenario will not be exactly equivalent to a clonally reproducing diploid population where the two chromosomes remain fully linked. In other words, a diploid population with independent segregation of chromosomes can be homozygous for a mutant allele, however a fully clonally propagating diploid population cannot be homozygous unless those mutations arose independently at the same site. Because tumor populations are diploid and fully clonal, i.e., do not undergo segregation of chromosomes, our diploid model is not directly applicable to them. However, a haploid asexual population with chromosomes of length 2*L* would accurately model and fully account for the dynamics in tumor populations (provided the SFS is constructed for 2*L* instead of *L* sites). In the case of viruses or other pathogenic organisms that reproduce asexually and do not recombine, the drug-resistant allele is known in many cases, e.g., in SARS CoV-2 (Moghadasi et al., 2023) and influenza (Behillil et al., 2020; Graitcer et al., 2011). Thus obtaining the SFS in the subpopulation carrying the drug-resistant allele can be informative of the strength of selection and timing of the selective sweep.

Presently, there are a few caveats of our study that would need to be addressed in order to apply them to natural or experimental populations. Firstly, we assume that the entire population has a constant size. It might be possible to incorporate a change in population size in this model in the future, allowing for a more general applicability of this model. This will be especially important in a tumor population, which itself is likely to be expanding. Secondly, we would likely require a large sample size to perform inference so that the crossover frequency can be observed in the sample SFS. Relatedly, the variance in the sample SFS from one replicate to another is larger for high frequency derived alleles, but much smaller for low frequency variants (Fig. S3), suggesting that in practice, identifying the *x*^−2^ decay at higher derived alleles could be challenging. As the purpose of this work was to study the population genetic quantities theoretically, the results presented here have been averaged over 10^6^ fixation events. However, an SFS obtained empirically usually represents a single replicate. Thus, the feasibility of accurate inference would have to be further explored. Thirdly, our theory works well when the beneficial mutation rate is not too high, i.e., when the dynamics of the population are not determined completely by multiple competing beneficial mutations (Gerrish and Lenski, 1998). The presence of pervasive clonal interference is difficult to determine empirically and might limit the application of the theory presented here to certain scenarios. With pervasive clonal interference, other theoretical frameworks like those proposed by Neher and Hallatschek (2013) and Melissa et al. (2022) will be more applicable.

## Acknowledgements

We thank Brian Charlesworth and Wolfgang Stephan for critically reading our manuscript and for providing helpful comments and suggestions that improved the manuscript. We thank three anonymous reviewers for providing helpful comments that improved the manuscript. We also thank Andrew Kern for helpful discussions about the project. This research was conducted using computational resources provided by ITS Research Computing at the University of North Carolina at Chapel Hill. This work was funded by the National Institutes of Health grant R35GM154969 to PJ.

### A Diffusion theory for changing population size

We first explain how the size *N* (*t*) of the subpopulation carrying the selected allele *A*^*∗*^ evolves (see Fig. 1). At the selected locus, we denote the three genotypes as *A*^*∗*^*A*^*∗*^, *A*^*∗*^*a*^*∗*^ and *a*^*∗*^*a*^*∗*^ with fitnesses 1 + *s*, 1 + *hs* and 1, respectively, where 0 *< h <* 1 is the dominance coefficient and 0 *< s <* 1 is the selection coefficient. In an infinitely large population, if the frequency of the selected mutant allele is *p* and that of wild-type allele is *q* = 1 − *p*, under Hardy-Weinberg equilibrium (HWE), the frequencies of the three genotypes are given by *p*^2^, 2*pq, q*^2^, respectively. We also assume that inbreeding occurs with probability 0 ≤ *F <* 1 so that the contribution to either homozygote’s frequency due to the union of heterozygotes is *Fpq*, and therefore, with inbreeding, the genotypic frequency vector in an infinitely large population is given by (Charlesworth and Charlesworth, 2010)

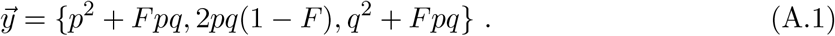

As expected, the frequencies are not in HWE due to non-random mating. The deterministic allele frequency *y*(*A*^*∗*^, *t*) = *y*(*A*^*∗*^*A*^*∗*^, *t*) + *y*(*A*^*∗*^*a*^*∗*^, *t*) of *A*^*∗*^ allele is then updated according to

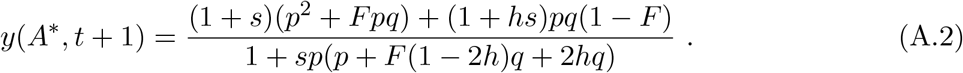

Then in the Wright-Fisher process, the number of mutants, *N* (*t* + 1) in generation *t* + 1 in a population of (census) size *N* are obtained from a binomial distribution with mean 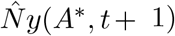,

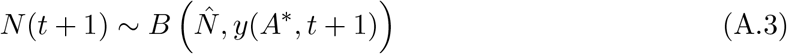

where 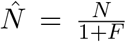 (Pollak, 1987; Glémin, 2012) to take care of the reduction in effective population size due to inbreeding. Note that while the population size *N* (*t*) is a binomially distributed random variable, as we are interested in the SFS when the selected allele sweeps, we require the distribution *F*_*c*_ of the allele frequency *conditioned* on fixation of the selected allele, an approximate theory for which has been developed for strong positive selection (Martin and Lambert, 2015; Götsch and Bürger, 2024).

We now consider the neutral population with alleles *a* and *A* whose number evolves according to a Wright-Fisher process with population size *N* (*t*) (see Fig. 1). If *n*_*a*_(*t*) and *n*_*A*_(*t*), respectively, denote the number of neutral wildtype and mutant individuals in generation *t* with *N* (*t*) = *n*_*a*_(*t*) + *n*_*A*_(*t*), the number of mutant individuals in the next generation is chosen according to a binomial distribution,

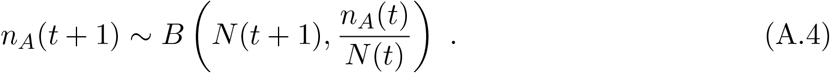

That is, the neutral mutants in the previous generation are sampled *N* (*t* + 1) times with equal probability.

To develop a diffusion theory for the neutral SFS in the changing population size for the model described above, we need to find the change in mean and variance of the neutral allele frequency, 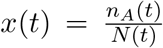 in one generation where *n*_*A*_(*t*) and *N* (*t*) are distributed according to Eq. (A.4) and *F*_*c*_, respectively. If … and 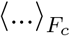 denote the average with respect to these respective distributions, it immediately follows from Eq. (A.4) that the mean neutral allele frequency at time *t* + 1 is given by

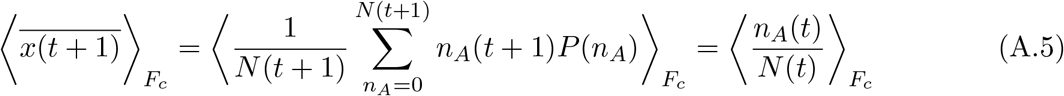

so that there is no change in the mean allele frequency. Similarly, the variance of the change in the allele frequency is given by

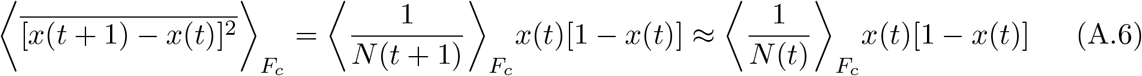

on using that 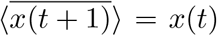. Thus, we find that the distribution of the mutant allele at neutral sites in a changing population obeys the following forward Fokker-Planck equation,

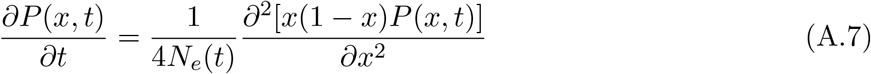

where, the variance effective population size (Waples, 2022)

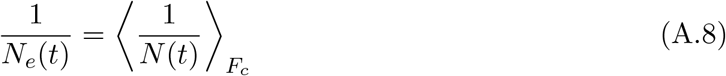

is the harmonic mean of the size of the subpopulation carrying selected allele conditioned on fixation.

On using Eq. (A.8) in Eq. (1) for the SFS, our numerics suggest that this approximation performs worse than if we assume that the size of the population varies deterministically. The reason is that while the above discussion on variance effective population size takes the fluctuations in *N* (*t*) due to strong but finite selection into account, it also implicitly assumes that the subpopulation sizes are completely uncorrelated in time which is possible only when the subpopulation size varies deterministically (Jain, 2025). To avoid the inconsistency in our assumptions, in this article, we assume that the size of the subpopulation changes according to Eq. (B.1) as described in Appendix B.

### Appendix B Deterministic dynamics of population size

If selection is strong (*Ns* 1) and the time at which the SFS is measured is not too small, we expect that the stochasticity in the population size can be ignored. Furthermore, the stochastic events leading to the eventual fixation of one copy of selected allele can be taken into account if the initial frequency of the selected allele is 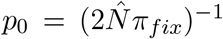 (Maynard Smith, 1976) where *π*_*fix*_ denotes the eventual fixation probability of a single mutant.

To this end, we first note that for small selection coefficient (*s* → 0), Eq. (A.2) shows that *y* ≈ *p* for any *F, h*, and in continuous time, it yields

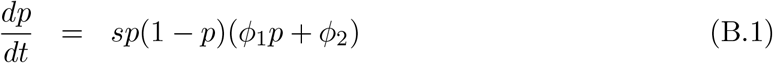

which gives

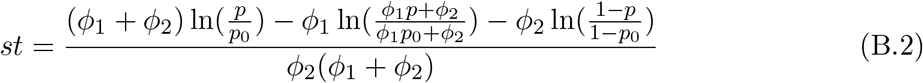

where,

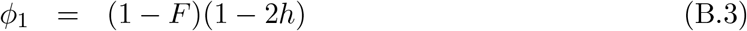

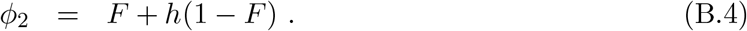

Thus, ignoring the fluctuations in the subpopulation size, we write

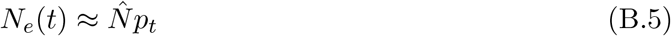

where the frequency *p*_*t*_ is determined from Eq. (B.2) with the initial condition, 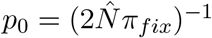 and for large *Ns*, the backward equation counterpart of Eq. (A.7) gives *π*_*fix*_ = 2*s*(*F* +*h*(1−*F*)) (Ewens, 2004).

### Appendix C Site frequency spectrum for changing population size

If the allele frequency at a neutral site in a population of changing size evolves according to Eq. (A.7) and is under strong selection so that 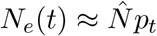 with *p* being the deterministically evolving allele frequency, the neutral SFS *f* (*x, t*) obeys Eq. (1) which can be solved exactly to give (Živković and Stephan, 2011; Jain and Kaushik, 2022)

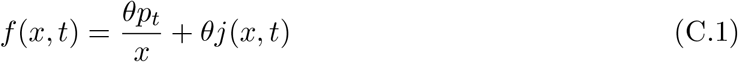

where,

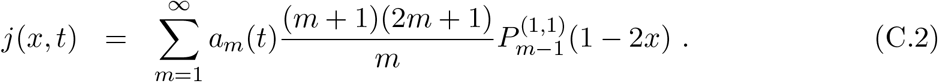

In the above expression, 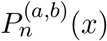 is the Jacobi polynomial and the time-dependent coefficients,

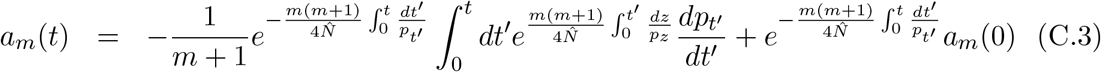

With

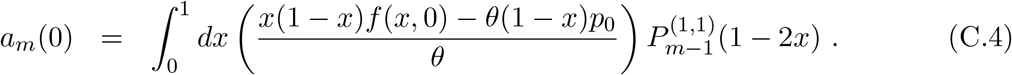

For the initial condition *f* (*x*, 0) = 0,

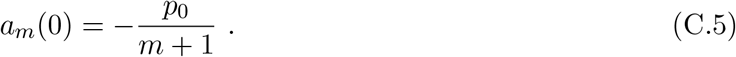

### Appendix D SFS during and after single selective sweep

As described in Appendix C, the exact expression for the neutral SFS in the infinites-sites model with changing population size 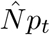 is given by Eq. (C.1). To our knowledge, the sum *j*(*x, t*) has not been previously analyzed for the SFS, and below we find an approximate expression for it when *p*_0_ → 0 and *p*_*t*_ → 1, that is, when the mutant is initially rare but has spread in the expanding population.

Due to Eq. (B.2), instead of time, it is convenient to express the SFS in terms of the allele frequency *p*_*t*_. Then

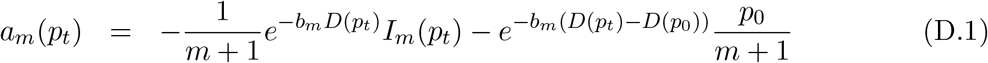

where,

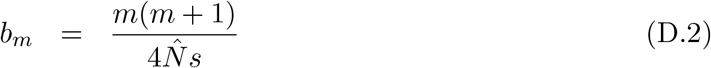

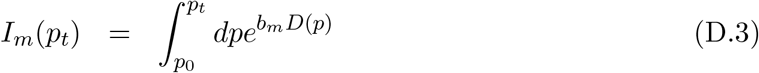

and due to Eq. (B.1),

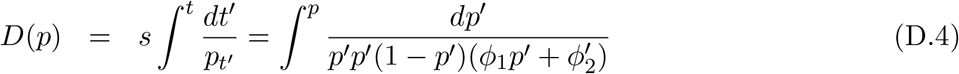

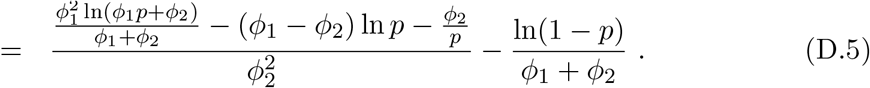

Since the population is expanding, *dD*/*dp* = (*pdp*/*dt*)^−1^ *>* 0, and the function *D* increases from 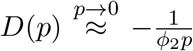 towards 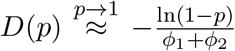 with increasing *p*. On expanding *D*(*p*) for small *p* and carrying out the integral in *I*_*m*_(*p*_*t*_), we obtain

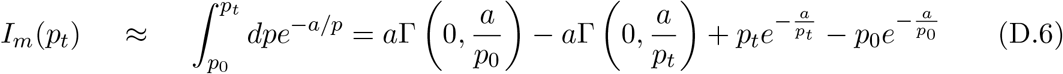

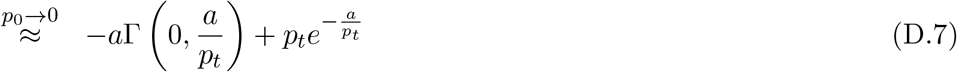

Where 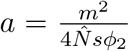. Also, for *p*_*t*_ → 1, as 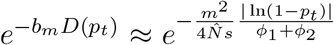 decays rapidly with *m*, the product 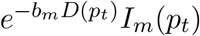 receives contribution from *I*_*m*_(*p*_*t*_) for small *a*/*p*_*t*_, and we can therefore write

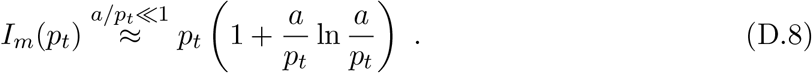

Furthermore, for large *m*, on approximating the Pochammer symbol 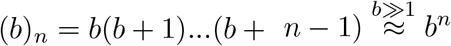, we can write

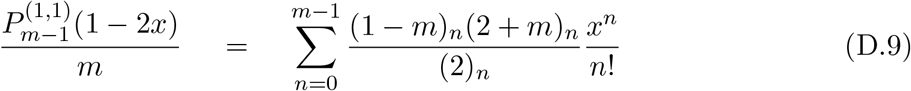

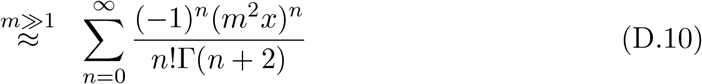

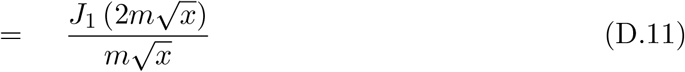

where *J*_*n*_(*x*) is Bessel function of the first kind.

Using the approximate expressions described above and approximating the sum over *m* on the RHS of Eq. (C.2) by an integral, we obtain

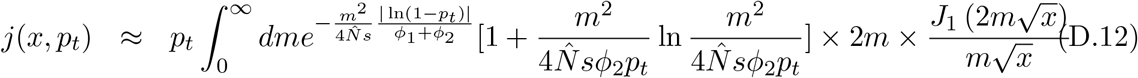

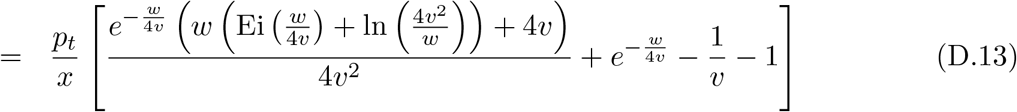

where 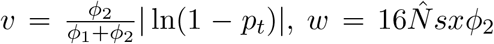, and 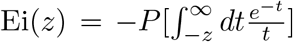 is the exponential integral function and *P* denotes the principal value of the integral. The constants φ_1_ and φ_2_ are given in Eq. (B.3) and Eq. (B.4), respectively.

---

**Figure S1.**
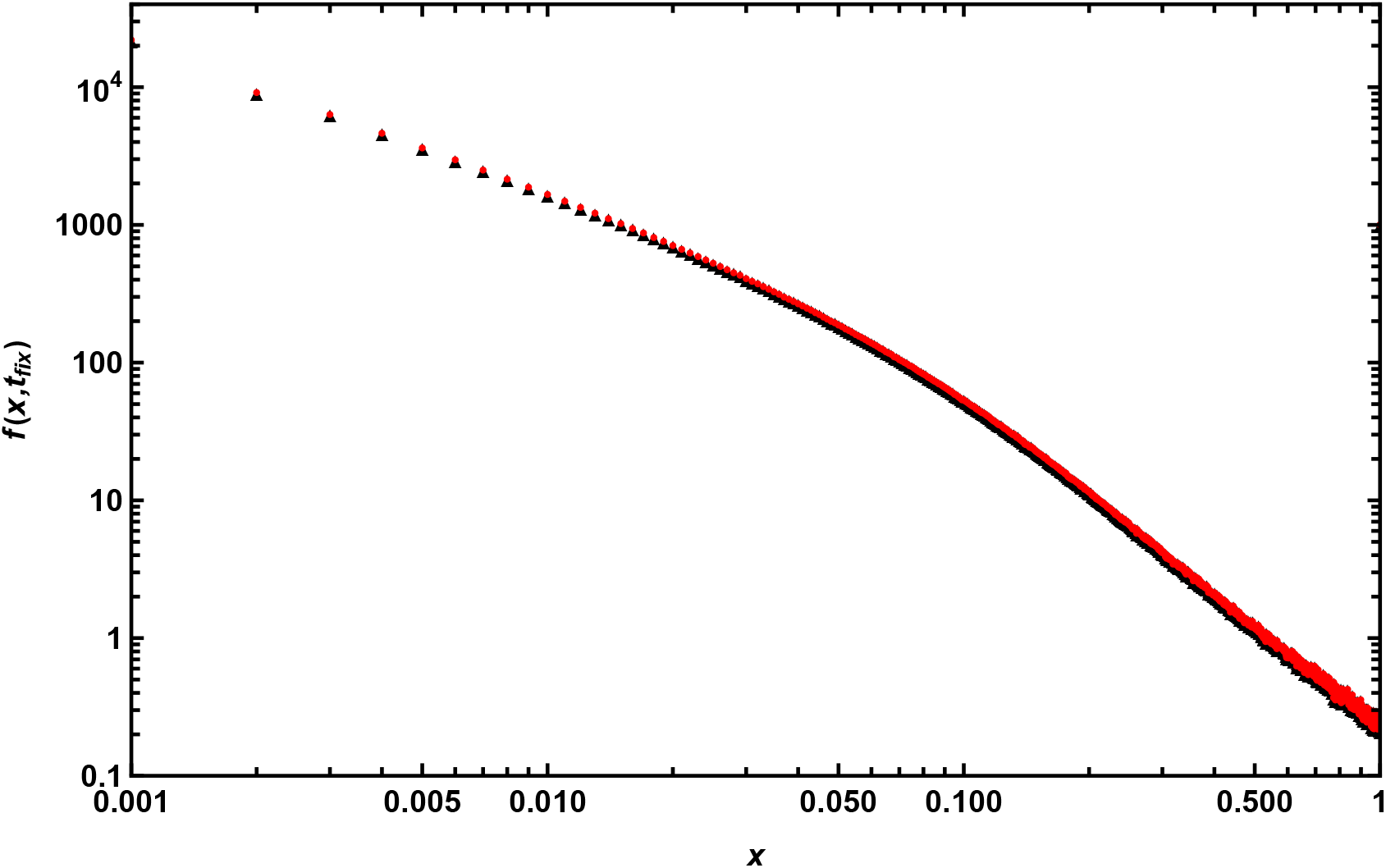
The SFS at neutral sites fully linked to a beneficial allele immediately post-fixation. Data is generated using forward-in-time simulations for *N* = 500 and *s* = 0.1. The red and black points correspond to two different initial conditions: when the neutral sites are at equilibrium (*θ*/*x*, black), and when the population is monomorphic (red) before the beneficial allele is introduced into the population. The parameters used are *µ* = 10^−7^, *L* = 10^5^, *F* = 0 and *h* = 0.5.

**Figure S2.**
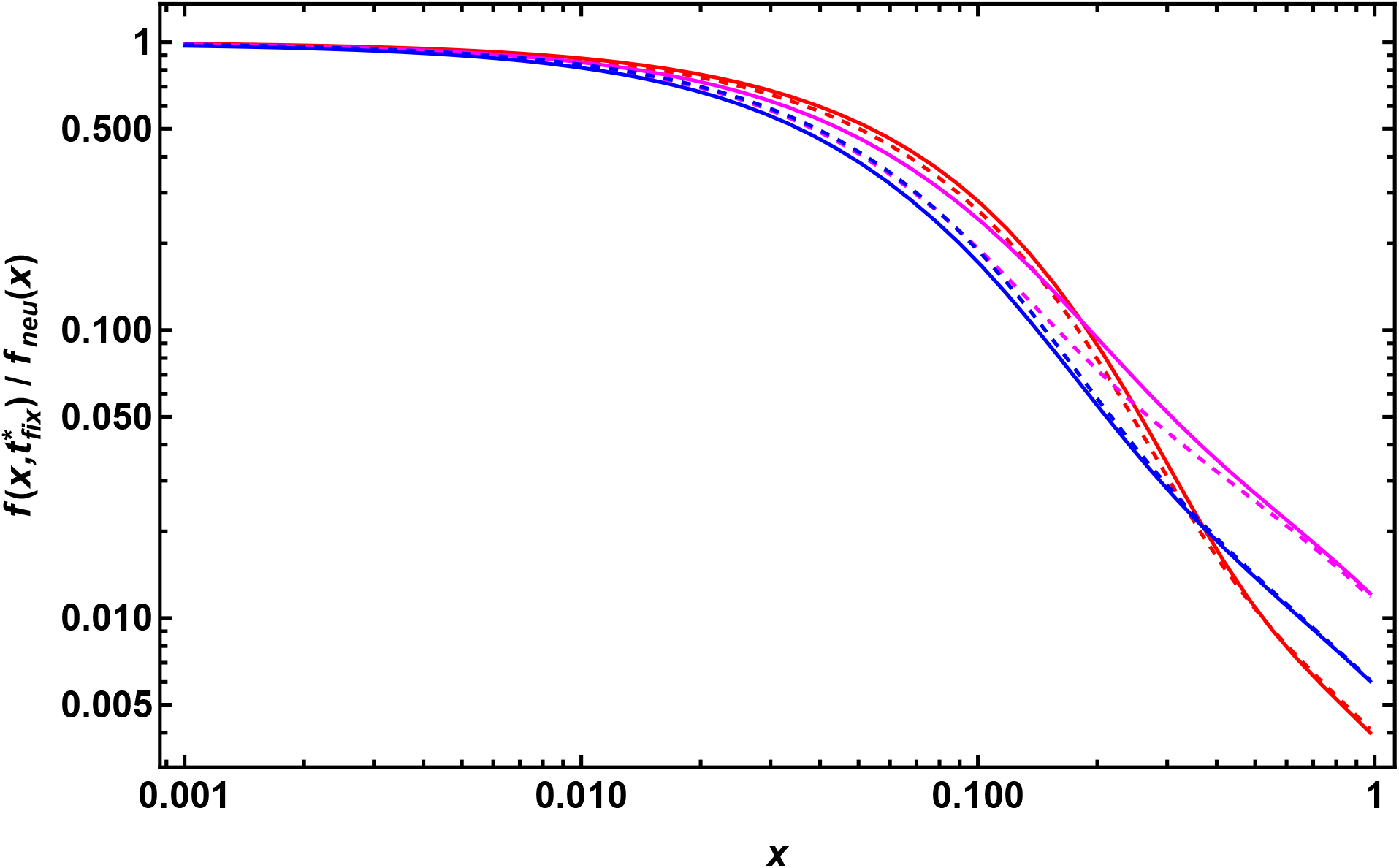
The SFS at neutral sites fully linked to a single new beneficial mutation immediately post-fixation relative to the neutral equilibrium SFS, *f*_*neu*_(*x*) = *θ*/*x*, for various dominance coefficients: a) *h* = 0.2 (magenta), b) *h* = 0.5 (blue), and c) *h* = 0.8 (red). The solid lines represent 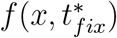, calculated at the conditional mean fixation time of the beneficial mutation [see Eq. (7)], while the dashed lines depict the SFS averaged over the entire conditional fixation time distribution. Both sets of lines are derived using diffusion theory. The parameters used are *N* = 10^3^, *µ* = 10^−7^, *s* = 0.1, *F* = 0, and *L* = 10^5^.

**Figure S3.**
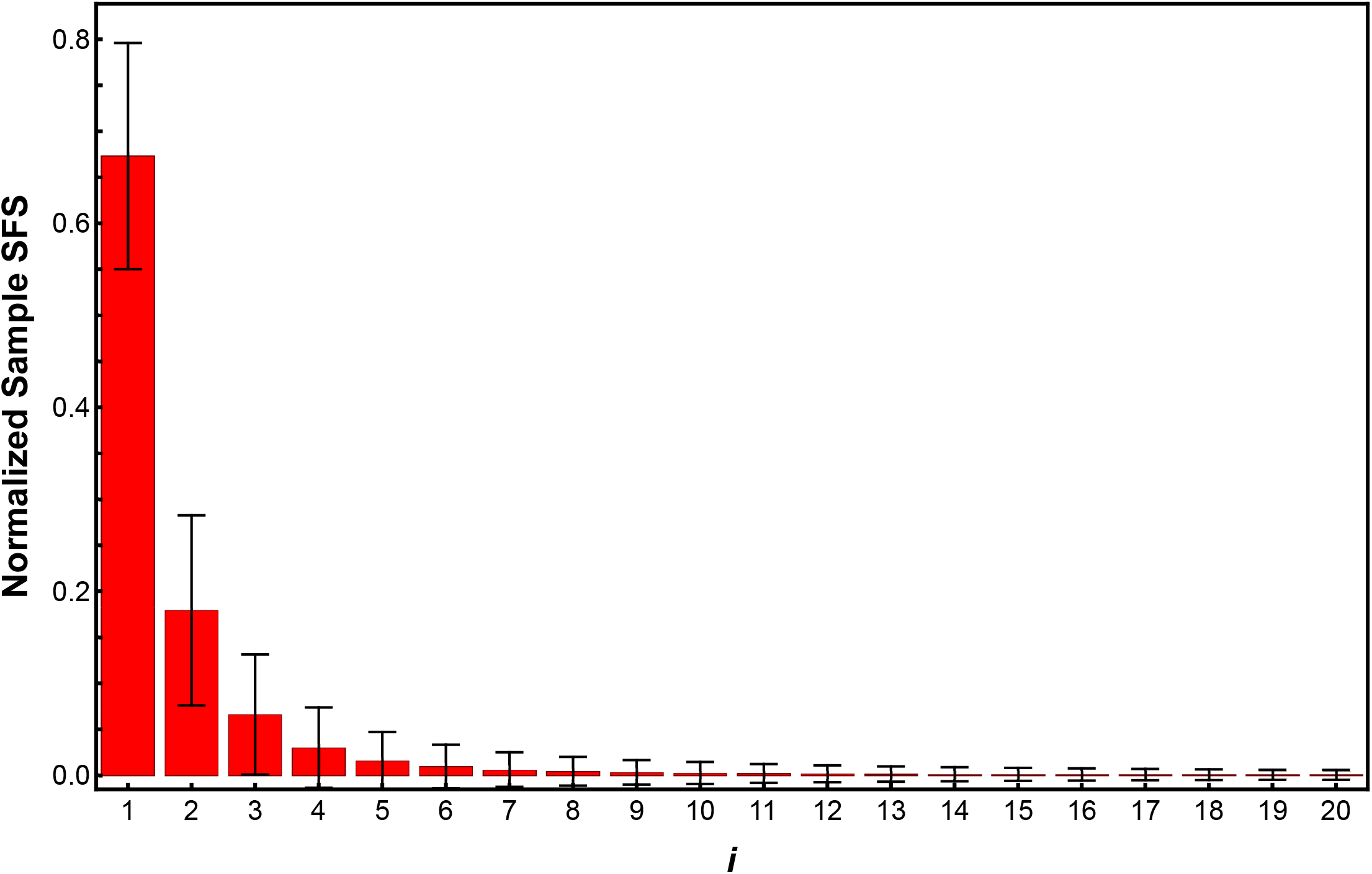
The normalized sample SFS at neutral sites immediately after the beneficial mutation reaches fixation. Here, the SFS is generated from 20 sampled genomes. The error bars represent standard deviation across 10^6^ independent simulation replicates. The parameters used are *N* = 10^3^, *µ* = 10^−7^, *s* = 0.1, *F* = 0, and *L* = 10^5^.

**Figure S4.**
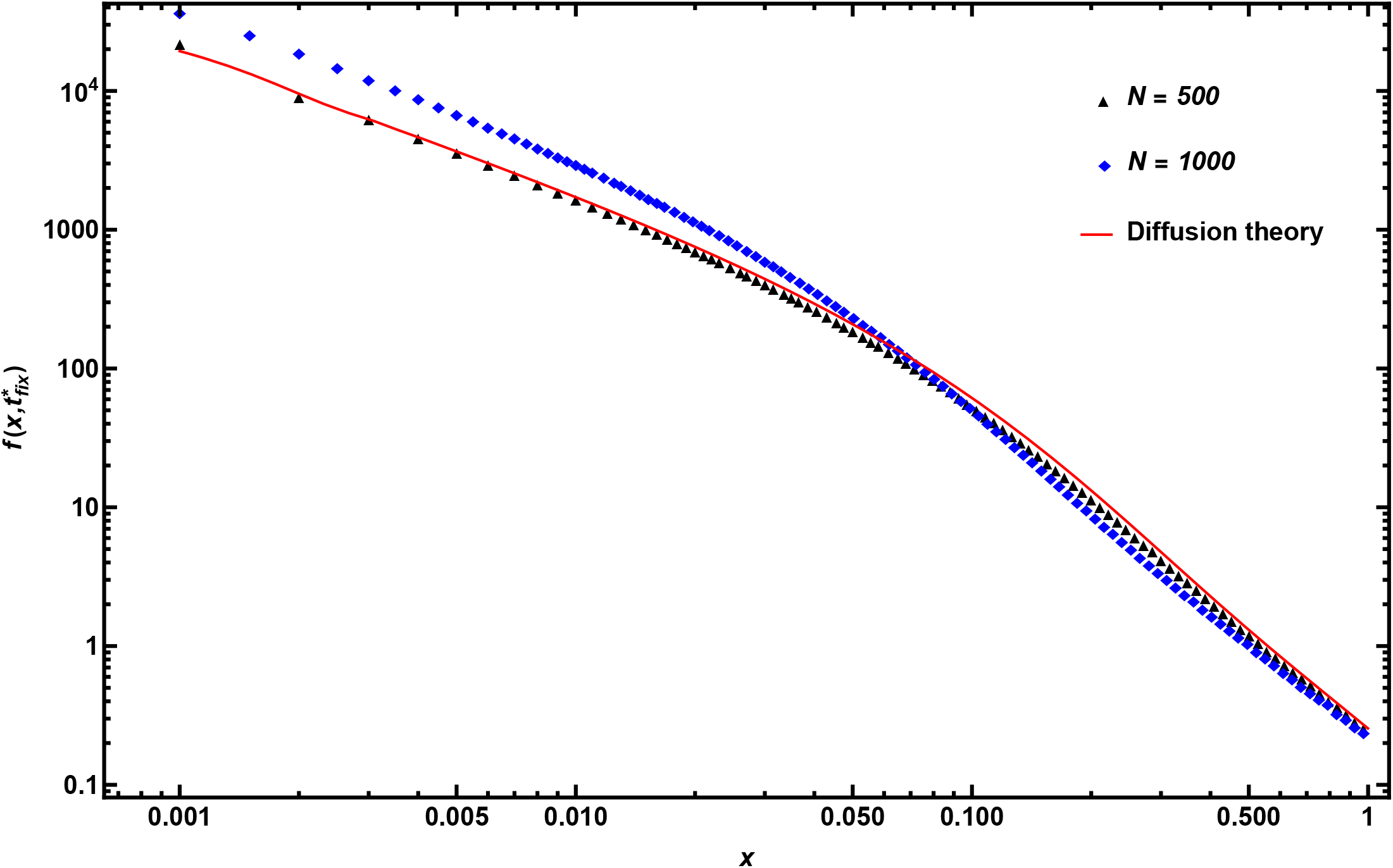
The SFS at neutral sites fully linked to a single new beneficial mutation immediately post-fixation is shown for varying population sizes: a) *N* = 500, and b)*N* = 1000, keeping the selection coefficient *s* fixed. The points are obtained using forward time simulations while the solid line is obtained using diffusion theory described by Eq. (1) for *N* = 500. The parameters used are *s* = 0.1, *µ* = 10^−7^, *L* = 10^5^, *F* = 0, and *h* = 0.5.

**Figure S5.**
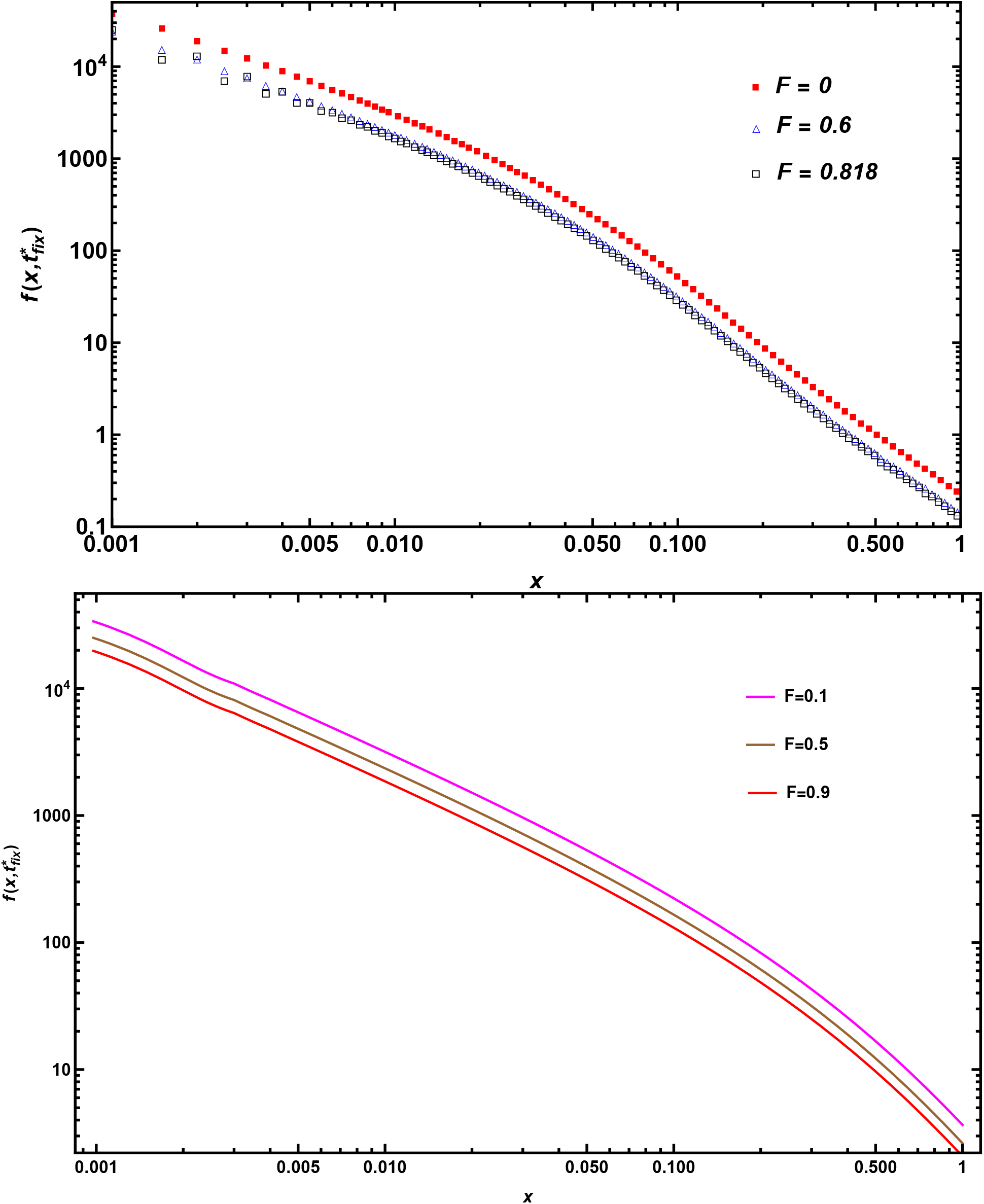
The SFS at neutral sites fully linked to a single new beneficial mutation immediately post-fixation is obtained from simulations (top) and diffusion theory (bottom) for various inbreeding coefficients. The other parameters used are *N* =10^3^, *µ* = 10^−6^ (top) and 10^−7^ (bottom), *s* = 0.1, *h* = 0.5 and *L* = 10^4^ (top) and 10^5^ (bottom).

**Figure S6.**
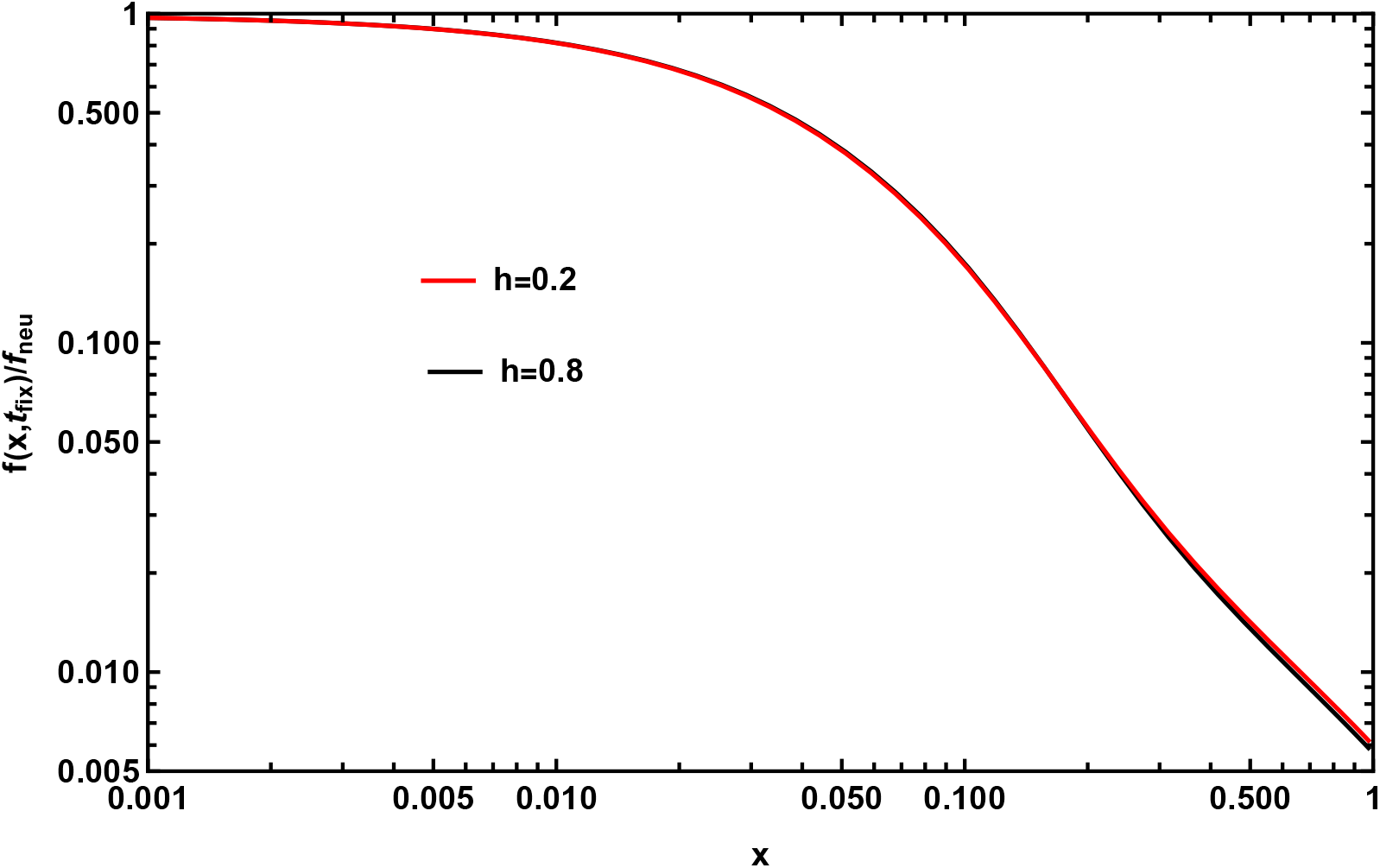
The SFS at neutral sites fully linked to a single new beneficial mutation following a selective sweep is shown for two different dominance coefficients: *h* = 0.2 and *h* = 0.8. The parameters used for this analysis are *N* = 10^3^, *µ* = 10^−6^, *s* = 0.1, *F* = 0.94, and *L* = 10^4^.

**Figure S7.**
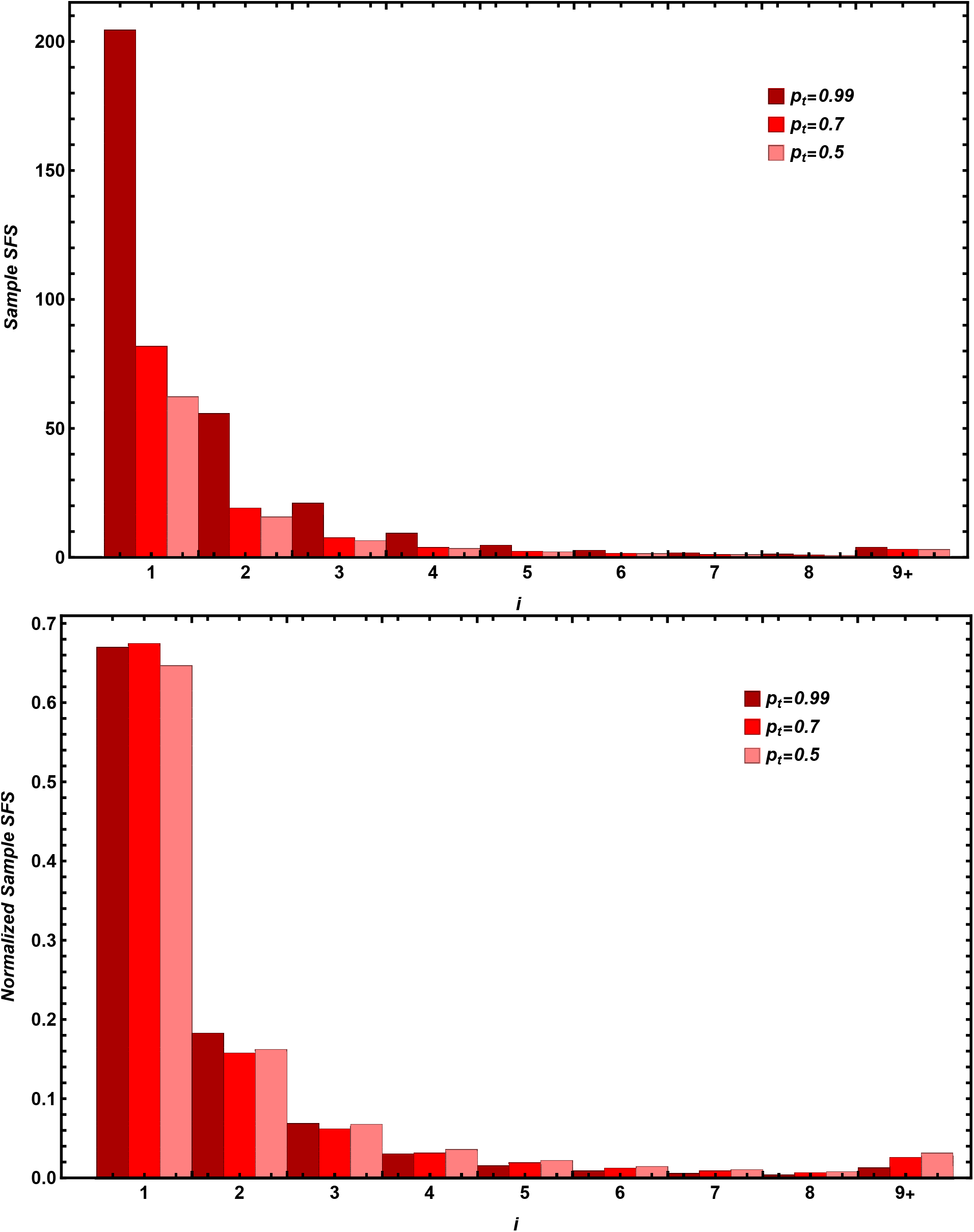
The sample SFS (above), and normalized sample SFS (below) at neutral sites when beneficial mutation frequency reaches the value mentioned in the legend. The data has been generated for 20 genomes from forward time simulations. The parameters used are *N* = 10^3^, *µ* = 10^−7^, *s* = 0.1, *F* = 0, and *L* = 10^6^.

**Figure S8.**
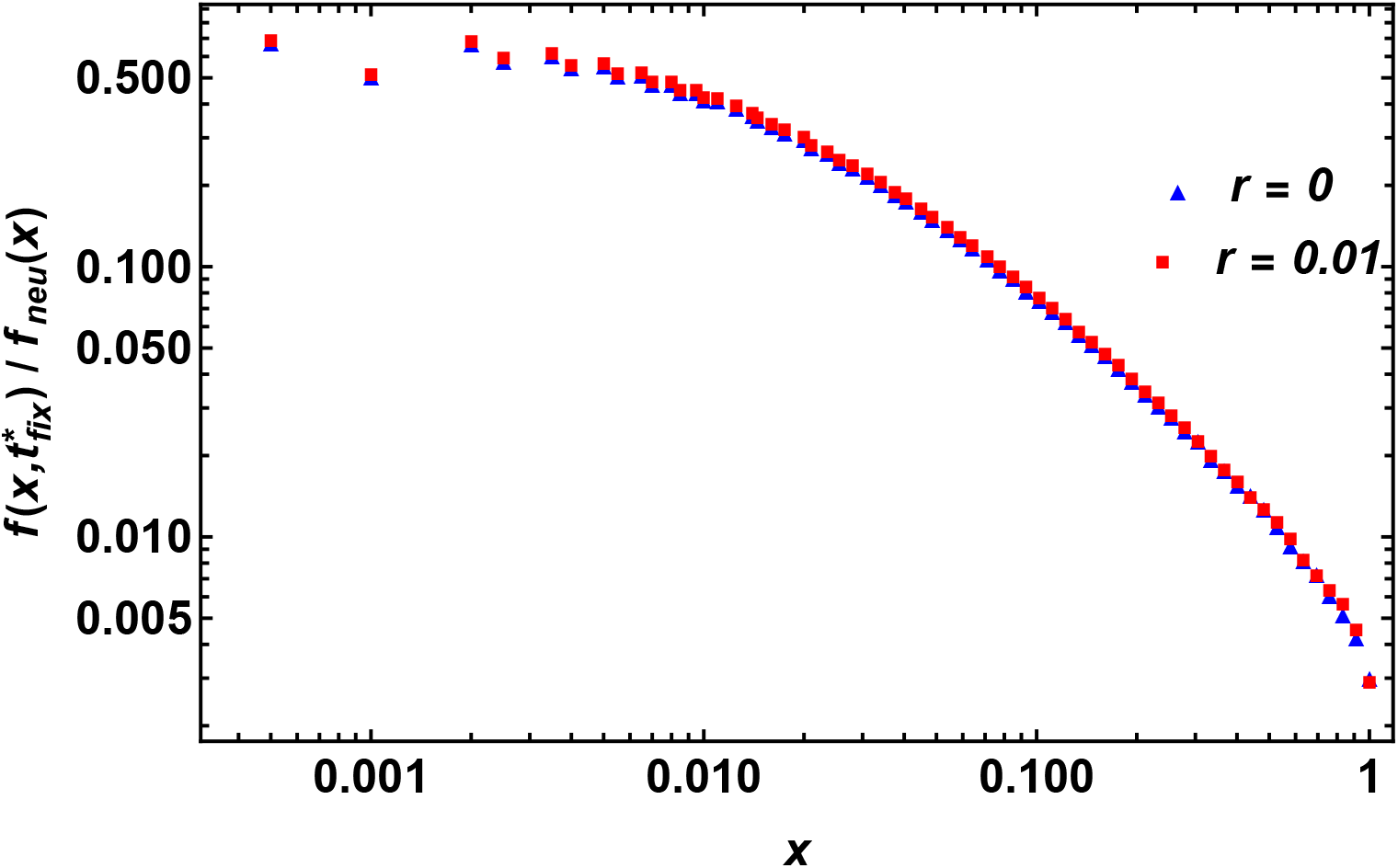
The figure shows the time-dependent SFS relative to the neutral equilibrium SFS, *f*_*neu*_(*x*) = *θ*/*x* in a strictly neutral expanding population with and without recombination; the simulation data are recorded when the deterministic allele frequency *p*_*t*_ = 0.35. Parameters used are *N* = 10^3^, *µ* = 10^−6^, *s* = 0.1, *L* = 10^4^, *F* = 0, and *h* = 0.5.

**Figure S9.**
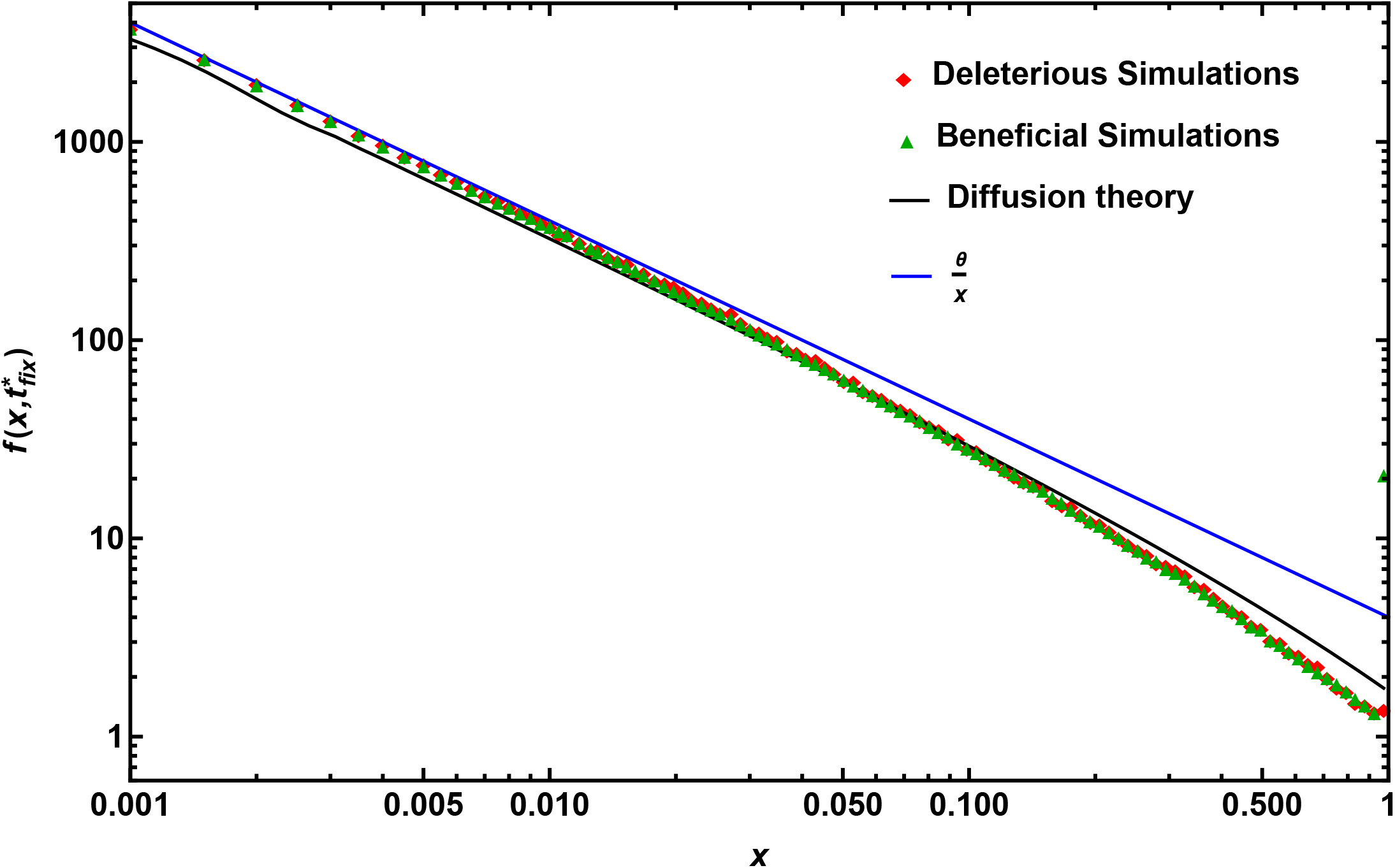
The simulation data for SFS at neutral sites when they are fully linked to a beneficial (green points) or a deleterious (red points) site immediately post-fixation are shown for *N* = 1000, and |*s*| = 0.002. The blue line shows the neutral equilibrium SFS, *θ*/*x*. The solid black line obtained by numerically solving Eq. (1) for the SFS in diffusion theory for beneficial hard sweep is shown for comparison and does not agree as well with the simulations results. The parameters used are *µ* = 10^−7^, *L* = 10^4^, *F* = 0 and *h* = 0.5.

**Figure S10.**
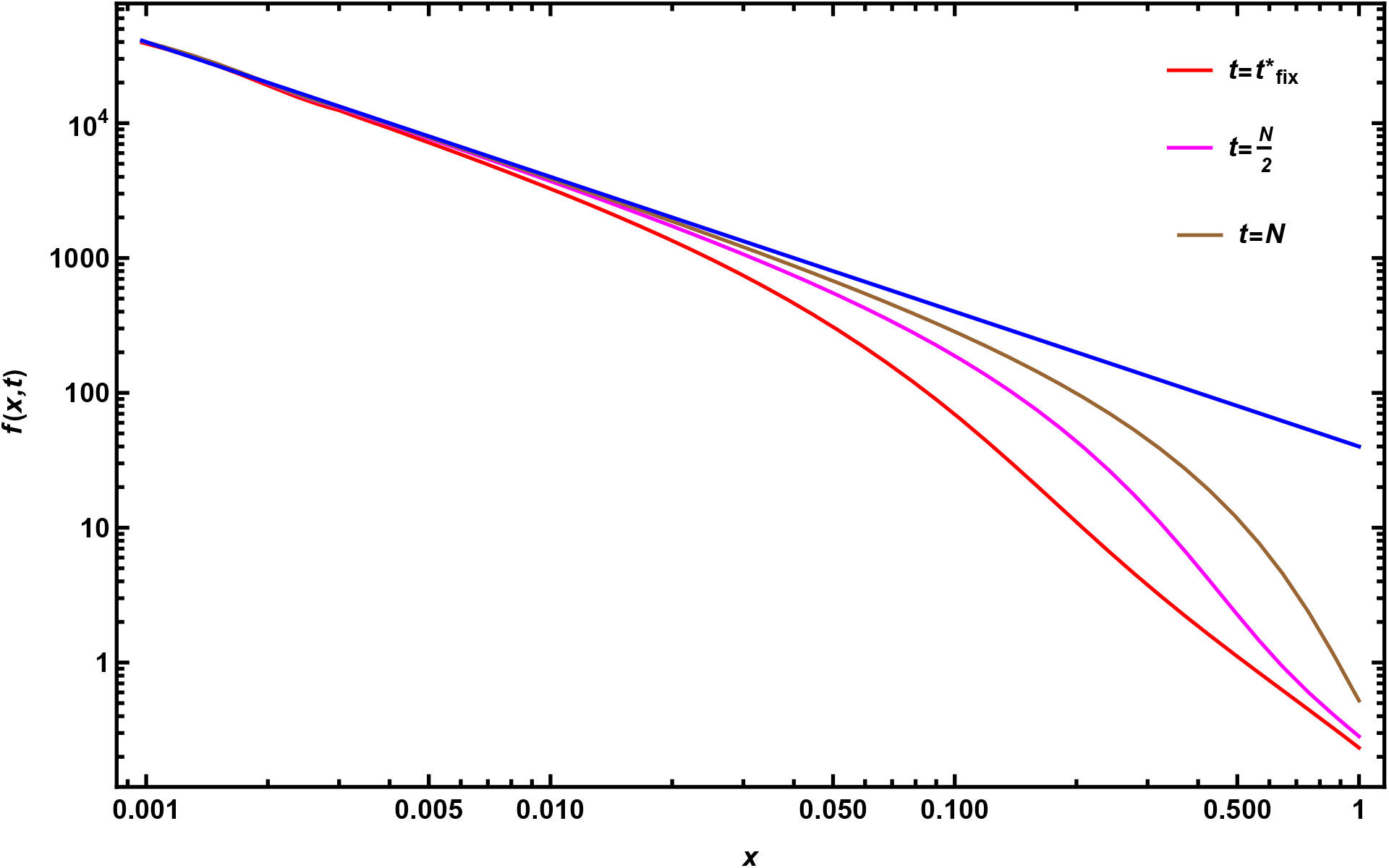
The recovery of the SFS at neutral sites fully linked to a single new beneficial mutation following a selective sweep is shown at different time points: *t* = *N* /2 (magenta) and *t* = *N* (brown). The SFS immediately post-fixation is depicted in red. Over time, the SFS gradually returns to the neutral equilibrium form, *θ*/*x* (blue). The parameters used for this analysis are *N* = 10^3^, *µ* = 10^−7^, *s* = 0.1, *h* = 0.5, *F* = 0, and *L* = 10^5^.

